# The encoding of touch by somatotopically aligned dorsal column subdivisions

**DOI:** 10.1101/2022.05.26.493601

**Authors:** Josef Turecek, Brendan P. Lehnert, David D. Ginty

## Abstract

The somatosensory system decodes a range of tactile stimuli to generate a coherent sense of touch. Discriminative touch of the body depends on signals conveyed from peripheral mechanoreceptors to the brain via the spinal cord dorsal column and its brainstem target the dorsal column nuclei (DCN)^1, 2^. Models of somatosensation emphasize that fast-conducting low- threshold mechanoreceptors (LTMRs) innervating the skin drive the DCN^3, 4^. However, post- synaptic dorsal column neurons (PSDCs) within the spinal cord dorsal horn also collect mechanoreceptor signals and form a second major input to the DCN^5–7^. The significance of PSDCs and their contributions to the coding of touch have remained unclear since their discovery. Here, we show that direct LTMR inputs to the DCN convey vibrotactile stimuli with high temporal precision, whereas PSDCs primarily encode touch onset and the intensity of sustained contact into the high force range. LTMR and PSDC signals topographically re-align in the DCN to preserve precise spatial detail. Different DCN neuron subtypes have specialized responses that are generated by unique combinations of LTMR and PSDC inputs. Thus, LTMR and PSDC subdivisions of the dorsal column encode different tactile features and differentially converge in the DCN to generate unique ascending sensory processing streams.

## MAIN TEXT

Fast-conducting low-threshold mechanoreceptors (Aβ-LTMRs) detect light mechanical forces acting on the skin and mediate discriminative touch^8–11^. Aβ-LTMR signals are conveyed rapidly from the periphery, and their axons ascend the dorsal column of the spinal cord and directly contact the dorsal column nuclei (DCN) of the brainstem. From the DCN, mechanosensory information is relayed to multiple targets in higher brain regions. Most sensory information is conveyed from the DCN to the somatosensory cortex via a prominent projection to the somatosensory ventral posterolateral thalamus (VPL) for the conscious perception of touch. A separate, lesser-known population of DCN neurons relays tactile information to the external cortex of the inferior colliculus (IC)^12, 13^ where it is integrated and contextualized with auditory information. Other populations of DCN neurons project to the olivocerebellar system^14, 15^ to coordinate motor adaptation, to secondary thalamic nuclei^16^ involved in affective state, and the spinal cord and periaqueductal gray^14, 16^. Thus, the DCN is a conduit of incoming mechanosensory signals, broadly connecting mechanoreceptors in the periphery to several major brain areas^17^.

Somatosensory coding in DCN neurons is known to be heterogeneous^18–21^, but how tactile signals are organized within the DCN and distributed to downstream targets remains unknown. Thus, we selectively recorded from DCN (gracile nucleus, **Extended Data Fig. 1**) neuron subtypes of the mouse using antidromic activation and optogenetic tagging to determine how sensory representations of the hindlimb are encoded at this early stage of the somatosensory hierarchy.

We found that different DCN neuron types encode distinct aspects of mechanosensory stimuli suited to their projection targets. VPL projection neurons (VPL-PNs) are the most abundant cell type in the DCN, outnumbering inferior colliculus projection neurons (IC-PNs) and local inhibitory interneurons (Vgat-INs) with an estimated proportion of VPL-PN:IC-PN:Vgat-IN of 2:1:1 (ref.^13^). We found that VPL-PNs had small excitatory receptive fields with large regions of surround suppression (**Fig. 1A-C, P-Q, Extended Data Table 1-2**). These VPL-PNs could entrain their firing to mechanical vibration, but for the vast majority this entrainment was restricted to a narrow range of frequencies below 150 Hz (**Fig. 1C-D**). DCN neurons projecting to the IC could be classified into several sub-groups (**Extended Data Fig. 2**), but most commonly had very large and exclusively excitatory receptive fields that included the entire hindlimb (**Fig. 1F-H, P**). Unlike VPL-PNs, most IC-PNs could entrain their firing to mechanical vibration across a broad range of frequencies (**Fig. 1H-I**). Vibration of the hindlimb evoked precisely timed action potentials that were entrained and phase-locked to vibrations ranging from 10 Hz up to 500 Hz, the highest frequency tested. Thus, neurons projecting to the VPL are tuned to convey finely detailed spatial information, whereas neurons projecting to the IC poorly encode spatial detail and are better suited to encode a broad range of mechanical vibrations that may be correlated with auditory stimuli. Vgat-INs had a wide range of receptive field sizes, lacked inhibitory surrounds, and unlike PNs typically lacked spontaneous firing (**Fig. 1K-N, P-R**). All three DCN cell types rapidly adapted to step indentations at low forces (**Fig. 1E, J, O**). In contrast to other cell types, VPL-PNs could more reliably encode the static phase of sustained high force indentation, well above forces at which Aβ-RA-LTMRs and Aβ-SA-LTMRs plateau (**Fig. 1E**)^22–25^.

**Figure 1:**
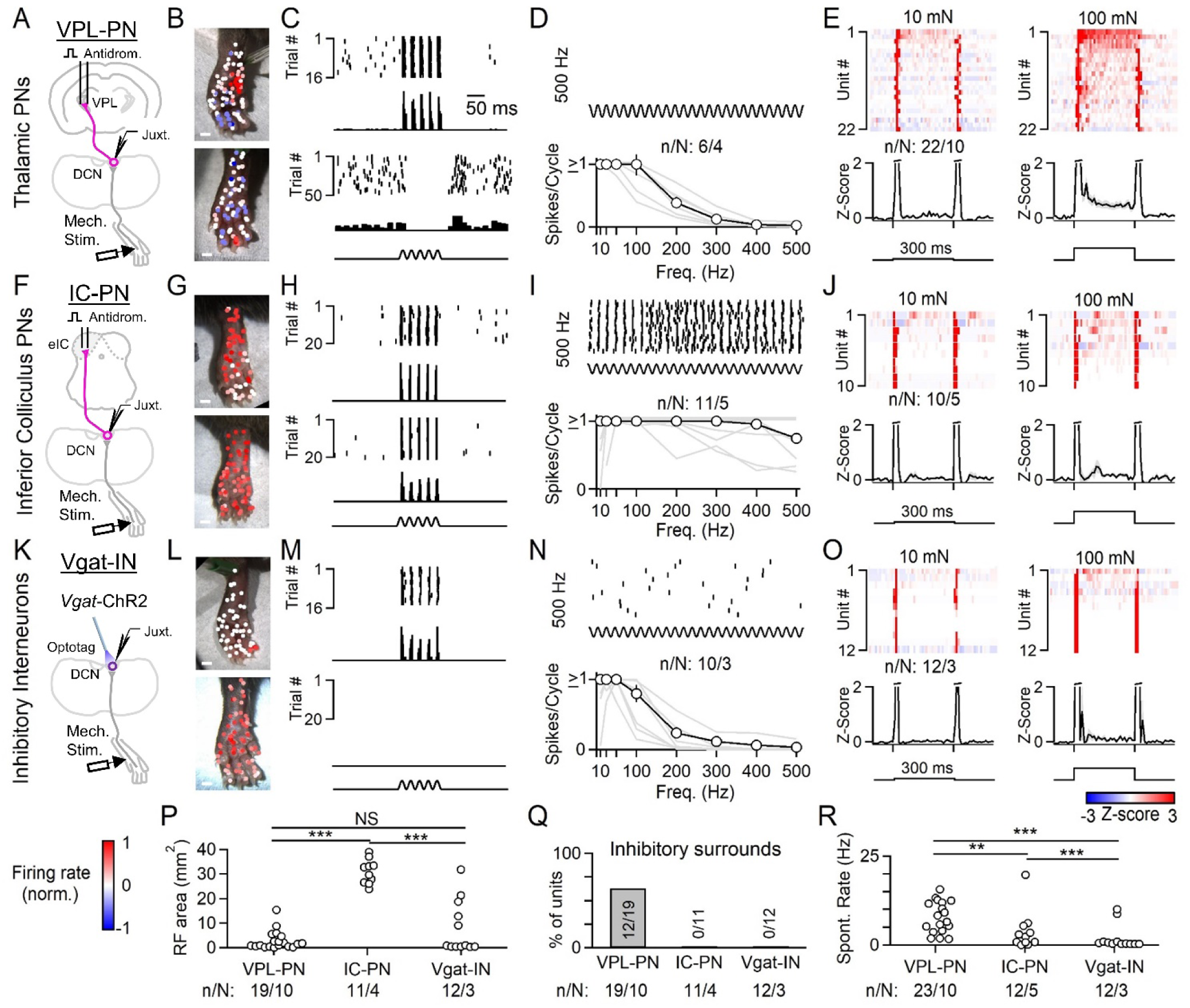
Tactile features are directed to distinct targets in the somatosensory hierarchy. **A.** Units in the DCN were recorded juxtacellularly in urethane-anesthetized mice. A stimulus electrode was inserted into the VPL and VPL projection neurons (VPL-PNs) were identified through antidromic activation and collision testing. The response properties of these units were then measured using a mechanical indenter with a 1 mm probe tip. **B.** Receptive fields of two different VPL-PNs. Each point is a single trial color coded by the unit’s normalized firing rate in response to a 100 ms 50 Hz vibration (10-20 mN). All scale bars 1 mm. **C.** Example trials from B with raster and histogram for two locations of the receptive field for the same unit: an excitatory receptive field (top) and inhibitory surround (bottom). Histograms for the excitatory RF are 1 ms bins, inhibitory surrounds 10 ms bins. **D.** Top: Example raster of VPL-PN response to a 500 Hz vibration delivered to the excitatory receptive field. Bottom: Vibration tuning of all VPL-PNs with average across units (black) and individual units (gray). **E.** Average responses (Z-scored firing rate) to step indentation in VPL-PN units (top) and average of all units (bottom) for 10 mN (left) and 100 mN (right). 300 ms indentations were delivered to the center of the excitatory receptive field using a 1 mm blunt-tipped probe. Histograms are in 10 ms bins. Average of all units shown at bottom as mean ± SEM. **F-J**. Same as A-E, but for units identified as projecting to the inferior colliculus (IC-PNs). **K-O**. Same as A-E, but for optotagged local inhibitory interneurons (Vgat-INs). **P.** Excitatory receptive field size area for all identified units. **Q.** Percentage of DCN cell types with detected inhibitory surrounds. **R.** Spontaneous firing rate of all units for each cell type. Number of experiments shown as n units in N animals (n/N). Statistical tests are K-S tests. Number of experiments and statistics also in Extended Data Table 1-2.

We next addressed how the unique response properties of VPL-PNs and IC-PNs are generated. Aβ-LTMR axons that travel via the dorsal column form the ‘direct’ dorsal column pathway from the skin to the DCN, and synapse directly upon DCN PNs and interneurons. However, post- synaptic dorsal column neurons (PSDCs) of the spinal cord, which receive input from a broad array of somatosensory neuron subtypes, including Aβ-LTMRs, also project to the DCN via an ‘indirect’ dorsal column pathway that exists across mammals, including primates^26–34^. PSDCs form up to 40% of the axons ascending the dorsal column (**Extended Data Fig. 3**)^35^, but the function of PSDCs and their contribution to somatosensory representations in the DCN and higher brain regions have remained unclear since their discovery. We hypothesized that direct and indirect dorsal column pathway projections contribute uniquely to the distinct tuning features of DCN neuron subtypes.

In order to isolate the functions of direct and indirect dorsal column inputs to DCN neuron responses, we used the light-activated chloride channel Acr^36^ to reversibly silence axon terminals of ascending inputs. We first generated *Cdx2-Cre; Rosa26^LSL-Acr^*^1^ mice to express Acr1 in all neurons below the neck to reversibly silence both primary sensory (direct pathway) and PSDC (indirect pathway) neurons that provide input to the DCN (**Fig. 2A; see methods**). We transiently silenced axon terminals in the DCN by preceding mechanical stimuli with brief, 300- 400 ms light ramps (**Extended Data Fig. 4**), which was optimal for suppressing excitatory inputs (**see Methods**). Mechanical stimuli were delivered in the final 100-200 ms of light application. Using this strategy to silence both the direct and indirect dorsal column pathway inputs abolished almost all DCN responses to vibration and low force step indentation of the hindlimb (**Fig. 2B- E**). We next selectively silenced all direct dorsal column pathway (Aβ-LTMR) input using *Avil^Cre^; Rosa26^LSL-Acr^*^1^ mice, allowing us to determine how the indirect pathway contributes to responses in individual DCN neurons (**Fig. 2F**). When light ramps were applied to silence Aβ- LTMR axon terminals in the DCN to block direct pathway inputs, the amplitude of responses was reduced but not eliminated. The indirect pathway was especially able to convey signals from low-frequency (10 Hz) mechanical stimuli to generate responses in the DCN (**Fig. 2G-H**).

**Figure 2:**
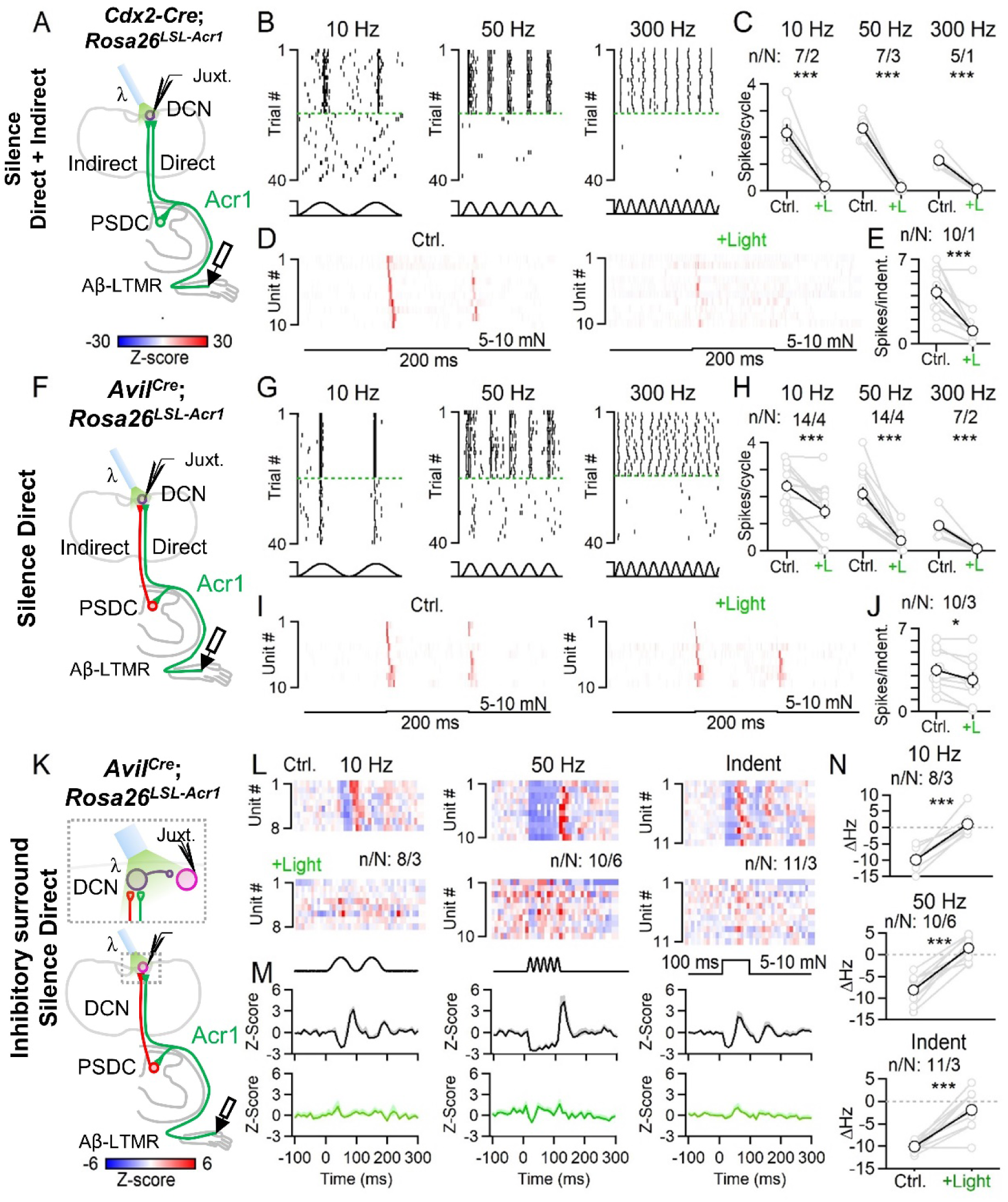
The direct dorsal column pathway conveys high-frequency vibration and fine spatial information to the DCN. **A.** In order to silence all ascending input to the DCN, Acr1 was expressed in both the direct and indirect pathway using *Cdx2-Cre; Rosa26^LSL-Acr^*^1^ mice. *Cdx2-Cre* drives recombination below C2 (ref.^53^); all spinal cord and DRG neurons below C2 express Acr1. Light was applied to terminals in the DCN to silence inputs in interleaved trials. 300-400 ms ramps of light (0.46 mW/mm^2^) preceded and overlapped with 100-200 ms mechanical stimuli. **B.** Rasters of single random DCN unit responses to 10 (left), 50 (middle) or 300 (right) Hz vibration under normal conditions or when silencing all ascending input. Silencing trials sorted and separated by dashed green lines. Vibrations were 10-20 mN. **C.** Average number of spikes per cycle during vibration under normal conditions or when silencing all ascending input for individual units (gray) and average across units (black). **D.** Average histograms (Z-scored firing rate) of responses to indentation (5-10 mN, 200 ms) under normal conditions (left) and when silencing all ascending input (right). **E.** Average number of spikes triggered by indentation at baseline, and when ascending input is silenced. **F.** In order to silence the direct pathway, Acr1 was selectively expressed in sensory neurons using *Avil^Cre^; Rosa26^LSL-Acr^*^1^ mice; remaining responses in DCN are mediated primarily by PSDCs. **G-J**. Same as B-E, but when silencing the direct pathway only. **K-N**. In order to determine the contribution of direct vs. indirect input to surround inhibition driven by local Vgat-INs, Acr1 was expressed in the direct pathway in *Avil^Cre^; Rosa26^LSL-Acr^*^1^ mice. Units in the DCN with inhibitory surrounds were identified, and the direct pathway was silenced during mechanical activation of the inhibitory surround. **L.** Average response of DCN units to stimulation of their inhibitory surrounds, evoked by 10 (left) and 50 Hz (middle) vibration, or indentation (right) under control conditions (top), or when silencing the direct pathway (bottom). **M.** Average inhibitory response across all DCN units in response to 10 and 50 Hz vibration, and indentation under normal conditions (top) and when silencing the direct pathway (bottom). **N.** Change in firing rate for each unit in response to stimulating the inhibitory surround under control conditions or when silencing the direct pathway. Summary plots are shown with individual units (gray), and average ± SEM (black). Number of experiments shown as n units in N animals (n/N). Statistical tests are paired t-tests. Number of experiments and statistics are also in Extended Data Table 1-2.

Silencing LTMR input also affected the response to step indentations, but the indirect pathway was able to reliably convey the onset and, in most units, the offset of low-threshold step indentation stimuli (**Fig. 2I-J**). On the other hand, indirect pathway input failed to evoke firing in response to 50 and 300 Hz vibratory stimuli. These findings were consistent across randomly sampled DCN neurons, as well as in identified VPL-PNs and IC-PNs, suggesting that the direct pathway drives high-frequency vibration across DCN cell types (**Extended Data Fig. 4**). Thus, high-frequency time-varying light touch stimuli such as vibration are exclusively encoded by the direct dorsal column pathway, whereas both the direct and indirect dorsal column pathway contribute to responses to gentle or light skin displacement across DCN neurons. The indirect pathway can also contribute to low-frequency vibration.

VPL-PNs can encode spatial information in part due to their prominent inhibitory surround receptive fields, and we therefore asked how the direct and indirect dorsal column pathways contribute to surround inhibition of these PNs (**Fig. 2K**). Spontaneously active VPL-PNs can be effectively inhibited by applying brief vibratory stimuli or indentation to areas outside of their excitatory receptive field. (**Fig. 2L, M**). Silencing Aβ-LTMR axon terminals in the DCN almost completely abolished inhibitory surrounds generated by 10 or 50 Hz vibratory stimuli and indentation (**Fig. 2L-N**). Thus, the direct dorsal column pathway is the primary driver of inhibitory surround receptive fields in VPL-PNs generated by these stimuli.

We next asked how PSDCs and the indirect pathway may contribute to DCN representations of other features of mechanical stimuli. As no genetic tools currently exist to selectively silence PSDCs, we used a pharmacological approach to block the indirect dorsal column pathway (**Fig. 3A**). Excitatory synaptic transmission in the lumbar spinal cord dorsal horn can be effectively blocked by application of glutamate receptor antagonists directly to the dorsal surface of the lumbar cord (**Extended Data Fig. 5**), thereby eliminating hindlimb-level PSDC neuron activation and contributions to responses recorded in the DCN. This glutamatergic transmission blockade was spatially restricted to the spinal cord region to which it was applied, since glutamate receptor antagonists applied to the lower thoracic cord did not affect DCN responses to mechanical stimulation of the hindlimb (**Extended Data Fig. 5**). When fast excitatory transmission was blocked in the lumbar spinal cord, DCN neurons could still entrain and phase- lock their spiking to a broad range of vibration frequencies of mechanical stimuli applied to the hindlimb (**Fig. 3B, C**). This finding, which is consistent with the Acr1 silencing experiments (**Fig. 2H**), indicates that Aβ-LTMR input via the direct dorsal column pathway underlies high- frequency vibration tuning in the DCN.

**Figure 3:**
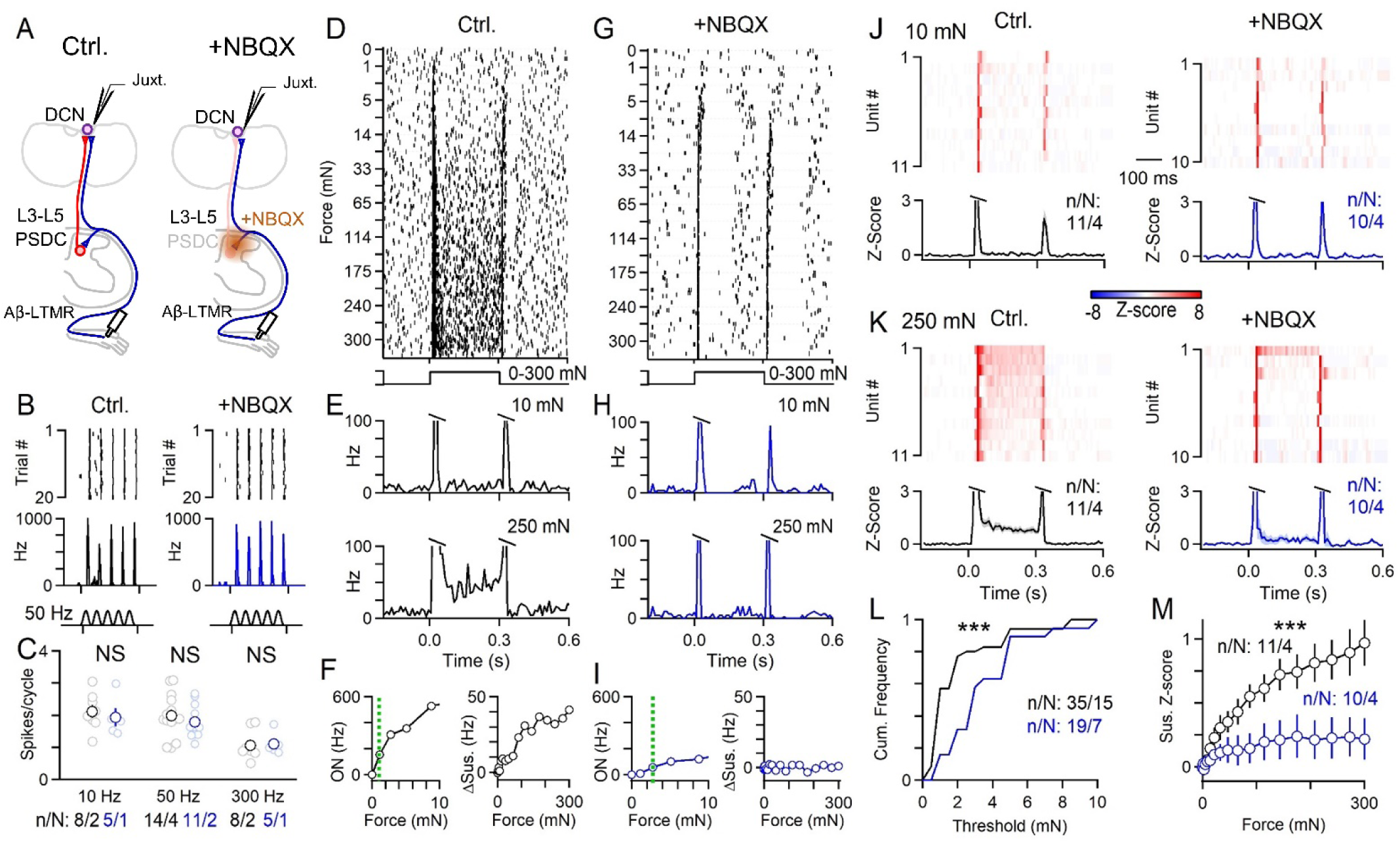
PSDCs and the indirect dorsal column pathway mediate wide dynamic range force intensity tuning in the DCN. **A.** Juxtacellular recordings made from random units (for vibration) or units with VPL-PN- like receptive fields (for indentation) in the DCN. An L3-L5 laminectomy and durotomy was performed to apply MK-801 (10 mM, 10 µL, 5 minutes) followed by NBQX (10 mM, 20 µL, for the recording duration). **B.** Example single unit response to 50 Hz vibration (10-15 mN) with raster (top) and histogram (bottom) of firing under control conditions (left), and a different unit after blockers were applied to the spinal cord (right). 1 ms bins. **C.** Summary of vibration responses for all units recorded under control conditions (black) or in the presence of blockers (blue). **D.** Raster of a single unit’s response to 300 ms step indentation at different forces (0-300 mN or 0-395 kPa) using a blunt probe (1 mm diameter) under control conditions. Trials originally interleaved, but sorted by force for presentation. High forces are innocuous in awake animals (see **Extended Data** Fig. 6). **E.** Histogram of firing for single unit in D-F for 10 mN (top) and 250 mN (bottom), 10 ms bins. **F.** Left: Max on response (0-20 ms) vs. indentation force for single unit in D. Dashed green line indicates threshold. Right: average sustained firing (100-300 ms) vs. force for single unit in D. **G-I**. Same as D-F, but for a different unit and in the presence of blockers. **J.** Average histograms for all units’ responses to 10 mN step indentation (top) and average across units (bottom) under control conditions (left). Responses to different DCN units in the presence of blockers in spinal cord (right). 10 ms bins. **K.** Same as J, but for 250 mN step indentation. **L.** Cumulative histogram of on response threshold for all units in control conditions (black) and for units in the presence of blockers (blue). **M.** Average sustained firing rate across units under control conditions (black) and for units in the presence of blockers (blue). Summary plots are shown with individual units (light), and average ± SEM (dark). Statistics are K-S tests. Number of experiments shown as n units in N animals (n/N). Number of experiments and statistics in Extended Data Table 1-2.

Another salient feature encoded by DCN neurons is stimulus intensity. We applied step indentations to the skin using blunt and smoothed probes (1 mm diameter) that generated graded, compressive stimuli from the low to high force range (1-300 mN). Although high forces were used, stimuli were applied over a wide area of skin. These stimuli were not noxious, as they failed to evoke paw withdraw or pain-related behavior in awake unrestrained animals, and most often generated no observable reaction (**Extended Data Fig. 6, Supplemental Video 1**). We found that under normal conditions DCN neurons were extremely sensitive to the onset and offset of low-force indentations, as expected, but also fired in response to sustained indentation of the skin, especially at high forces (**Fig. 3D-F**). High-force responses were prominent in VPL- PNs, and to a lesser extent in IC-PNs (**Fig. 1E, J**). Sustained firing during maintained step indentations was proportional to the force applied. DCN neurons could therefore not only detect the onset and offset of very gentle stimuli, but could also encode sustained mechanical stimuli across a broad range of intensities, into the high force range (**Fig. 3F**).

To determine the contribution of PSDCs to innocuous high-intensity stimuli, we recorded from DCN units that had receptive fields and spontaneous firing characteristic of VPL-PNs, the most abundant cell type in the DCN. When excitatory transmission was blocked in the spinal cord to suppress PSDC input, firing during the sustained phase of indentation was almost eliminated across all forces for most units (**Fig. 3G-K**). Thus, in the absence of PSDC input, DCN neurons could no longer encode the intensity of maintained stimuli (**Fig. 3M**). We also found that attenuating PSDC input to the DCN raised the threshold of DCN neurons to the onset of step indentation (**Fig. 3L**). These findings suggest that PSDCs provide graded force information to the DCN, enabling responses to sustained high intensity stimuli. At the same time, PSDCs also contribute to detection of gentle touch stimuli. Thus, PSDCs and the indirect dorsal column pathway are required for the wide dynamic range of intensity tuning in DCN neurons, enabling detection and encoding of a broad range of stimulus intensities.

We also found that the graded coding of intense stimuli in the DCN was relayed upstream to middle VPL (mVPL) neurons. Both low-threshold sensitivity and sustained responses to high-threshold stimuli in the mVPL were strongly dependent on the DCN (**Extended Data Fig. 7**). Lesioning the DCN also altered spontaneous firing in the mVPL (**Extended Data Fig. 7**), although this manipulation will also affect the corticospinal tract. Many neurons in the DCN are spontaneously active, but we found that lesioning the dorsal column did not have major effects on spontaneous firing in randomly recorded DCN neurons (n/N: 16/2), suggesting that PSDCs do not drive spontaneous firing in the DCN.

The observation that DCN neurons encode high intensity indentation stimuli is noteworthy because most Aβ-LTMRs are thought to saturate their firing at relatively low indentation forces^22–25^. As most of the high force responses of DCN neurons during sustained indentations are mediated by PSDC neurons, we considered the possibility that PSDCs transmit signals emanating from both LTMRs and high-threshold mechanoreceptors (HTMRs), which typically do not project directly to the DCN via the direct dorsal column pathway^37^. In order to activate HTMRs without concurrent activation of LTMRs, we expressed the light-activated cation channel ReaChR in somatosensory neurons expressing *Calca* (CGRP) using *Calca*-*FlpE*; *Rosa26^FSF-ReaChR^* mice^38, 39^ and applied light directly to the skin of the hindlimb (**Fig. 4A**). Among sensory neurons expressing *Calca* are A-fiber HTMRs, but also C-fiber HTMRs, thermoreceptors, and polymodal C-fiber neurons^40–43^. As before, VPL-PNs responded rapidly to strong but innocuous mechanical step indentations of the skin with additional long latency spikes (**Fig. 4B, D; Extended Data Fig. 8**). Optical excitation of *Calca*^+^ HTMRs within the same area of skin triggered firing in VPL-PNs with fast latencies that were consistent with A-fiber activation, but were longer than those evoked by mechanical stimuli (**Fig. 4B-E**). These rapid optogenetically-evoked responses were sometimes followed by a second and much longer latency burst (**Fig. 4E**). These two temporal components of firing were consistent with the activation of intermediate and slow conducting Aδ and C fiber neurons known to express *Calca*^41^. Strikingly, almost all VPL-PNs fired robustly and consistently to optical activation of *Calca*^+^ endings in the skin (**Fig. 4C, E**). The slow latency of these responses was likely not due to slow opsin kinetics, as activation of all endings in the skin, including Aβ-LTMRs, generated faster latency responses (**Extended Data Fig. 8**). Previous work has shown sparse *Calca*^+^ neuron input to the DCN^44, 45^. We observed very few fibers in the DCN labeled by *Calca*-*FlpE*, and direct optical stimulation over the DCN failed to evoke firing in the same units that could be activated by optical stimulation of endings in the skin (**Extended Data Fig. 8**). Moreover, responses in the DCN evoked by activation of *Calca*^+^ endings in the skin were dependent on synaptic transmission in the spinal cord, as they were strongly attenuated by addition of NBQX to the spinal cord (**Extended Data Fig. 8**). Responses in the DCN were mediated by the dorsal column, as severing the dorsal column eliminated light-evoked responses in the DCN (**Extended Data Fig. 8**). These findings suggest that high-threshold information is relayed to the DCN by PSDCs that receive input from HTMRs, either directly or through local interneuron circuits within the spinal cord dorsal horn.

**Figure 4:**
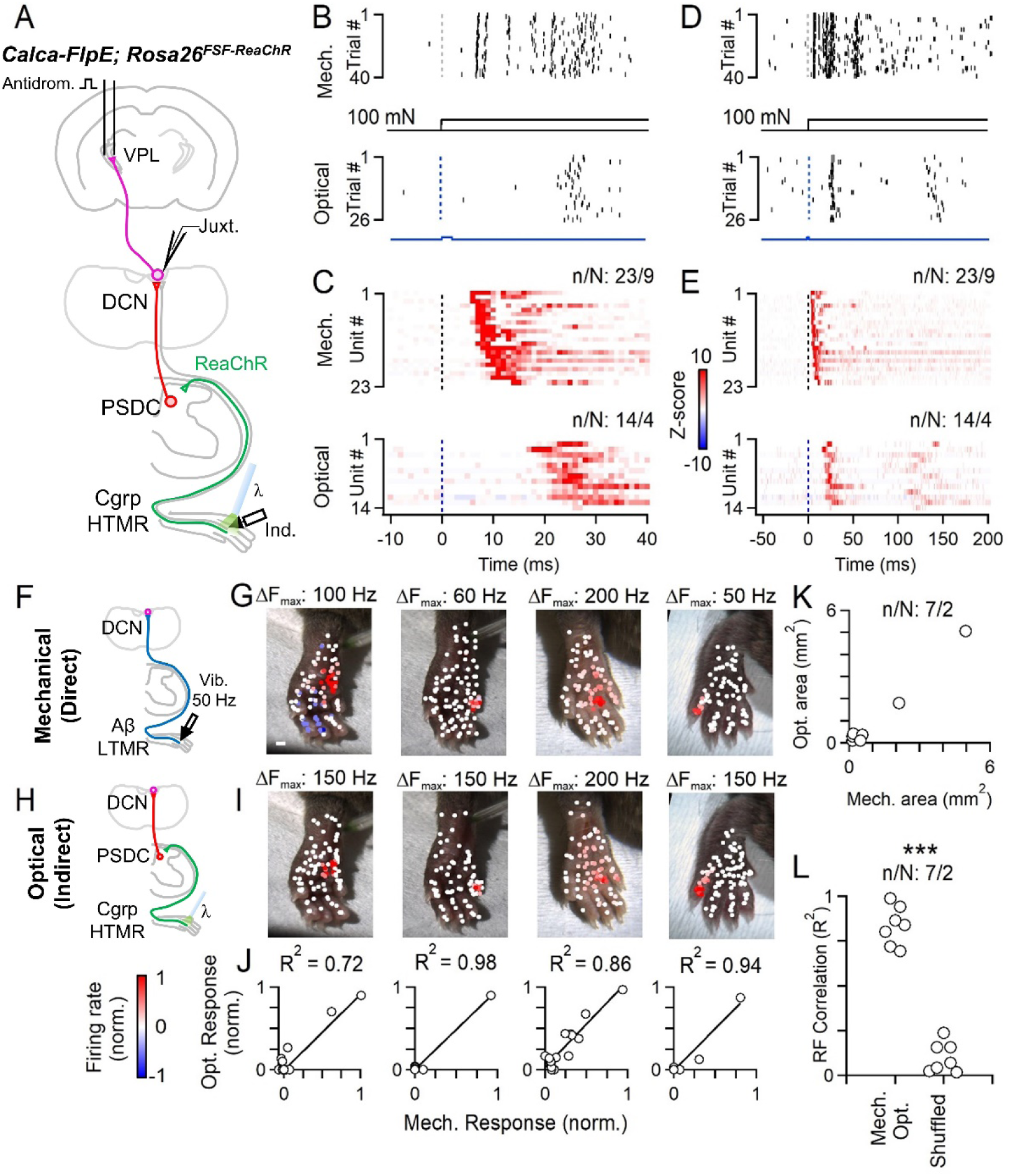
Somatotopic convergence of the direct and indirect dorsal column pathways onto individual VPL-PNs. **A.** VPL-PNs were identified with antidromic activation in *Calca*-*FlpE*; *Rosa26^FSF-ReaChR^* animals. Pulses of light activated ReaChR in *Calca* HTMRs in the hindlimb. **B.** Raster of a single VPL-PN unit response to indentation (100 mN, 1 mm probe tip, top), and the same unit’s response to optical activation (2 ms, 60 mW/mm^2^) of HTMRs in the skin (bottom). **C.** Histograms for all identified VPL-PN responses to mechanical indentation (100 mN, top), and other VPL-PN responses to optical activation of HTMRs in the skin (bottom). 1 ms bins. Z-score colorscale at right. **D-E**. Same as B-C, but longer timescale. **F.** A 50 Hz 100 ms vibration (10-20 mN) was applied to different points on the skin for putative VPL-PNs in *Calca*-*FlpE*; *Rosa26*^FSF-ReaChR^ animals. This stimulus activates the direct pathway. **G.** Vibration receptive field maps for four VPL-PNs. Each point is a single trial color coded by its normalized mechanically evoked firing rate, colorscale at bottom left. **H.** For same units in G, light pulses (5 pulses at 0.5 Hz, 2 ms, 60 mW/mm^2^) were applied to different points on the skin. This stimulus activates the indirect pathway. **I.** Optical receptive field for same units in G. Each point is a single trial color coded by its normalized optically evoked firing rate. **J.** Correlation of optical and mechanical responses. Each point is the average mechanically and optically evoked responses for a single region of skin. **K.** Area of optically evoked receptive field vs. area of mechanically evoked receptive field for single units. Each point is one DCN unit. **L.** Correlation coefficients (R^2^) of optically- and mechanically-evoked receptive fields within individual units, or R^2^ of receptive fields shuffled between units. Number of experiments shown as n units in N animals (n/N). Statistics in L are K-S tests. Number of experiments and statistics in Extended Data Table 1-2.

We also observed responses in IC-PNs evoked by stimulating *Calca*^+^ endings in the skin, but they were weaker than those seen in VPL-PNs and were restricted to a subset of the receptive field more sensitive to low-frequency mechanical stimuli (**Extended Data Fig. 9**). Activation of *Calca*^+^ endings failed to evoke inhibitory surrounds (**Extended Data Fig. 9**), consistent with the absence of sustained high force responses in Vgat-INs (**Fig. 1O**).

How are receptive fields of VPL-PNs shaped by the direct and indirect dorsal column pathway inputs? We measured the contributions of the direct and indirect dorsal column pathways to receptive fields of individual VPL-PNs. To do this, we determined Aβ-LTMR input contributions to VPL-PN receptive fields by using vibratory stimuli, as high-frequency vibration is encoded by the direct pathway (**Fig. 4F-G**). In the same experiment, we determined PSDC input contributions to receptive fields of the same VPL-PN neurons using optical activation of *Calca*^+^ endings in the skin, because *Calca*^+^ neuron inputs to the DCN are conveyed solely by the indirect pathway (**Fig. 4H-I**). Strikingly, the receptive fields of the direct and indirect dorsal column pathway inputs onto VPL-PNs were highly aligned (**Fig. 4G, I, J-L**). Responses to gentle vibration were typically restricted to a single digit or 1-2 pads, and optical activation of HTMRs evoked firing only in areas that were sensitive to vibratory stimuli, restricted to the same single digit or 1-2 pads. These findings suggest an elaborate somatotopic alignment of the periphery, spinal cord, and DCN: Aβ-LTMRs and HTMRs innervating a small area of skin project into the central nervous system and diverge, with Aβ-LTMRs projecting via the dorsal column directly to the DCN, and both LTMRs and HTMRs activating PSDCs in the spinal cord dorsal horn. Aβ-LTMRs (direct pathway) and PSDCs (indirect pathway) with similar receptive fields then re-converge within the DCN to enable representation of a broad array of tactile features for a single, small area of skin.

## DISCUSSION

The dorsal column system enables the perception of a rich array of tactile features^2–4, 11^. Models of discriminative touch have primarily focused on the roles of LTMR subtypes and their direct dorsal column pathway projections in the creation of these representations^4, 9, 11^. Here, we demonstrate that PSDCs are a critical component of a brainstem circuit that transforms ascending tactile inputs to produce specialized tactile representations.

Our findings support a new model of the dorsal column discriminative touch pathway **(Extended Data Fig. 10)**. Aβ-LTMRs and the direct dorsal column pathway underlie vibration tuning, whereas PSDC neurons and the indirect dorsal column convey the intensity of sustained stimuli. Both pathways detect the onset of stimuli and low-threshold responses, together providing exquisite sensitivity. Remarkably, these two components of the dorsal column pathway differentially converge upon distinct DCN-PN subtypes to generate unique combinations of response properties in different sensory streams. Information conveyed to S1 via VPL-PNs emphasize spatial detail, moderate-low frequency vibration (<150 Hz), and sustained stimulus intensity. These features are likely generated by prominent input from both Aβ-LTMRs and PSDCs. Tactile signals conveyed to the IC by IC-PNs encode broadband vibratory information (10-500 Hz) rather than spatial detail, and are likely mainly driven by LTMRs, especially Meissner corpuscle/Aβ RA1-LTMRs and Pacinian corpuscle/Aβ RA2-LTMRs. We found that LTMR input generates much of the surround inhibition in VPL-PNs, and that Vgat-INs do not fire in response to sustained high-force stimuli, suggesting that they do not receive HTMR input via PSDCs. We also found that PSDCs do not play a major role in surround inhibition, but it remains possible that they are involved in other aspects of mechanically-evoked or tonic inhibition that we did not measure. Many of the DCN response properties reported here can be observed in both anesthetized and non-anaesthetized conditions^46–48^, but PSDCs may have additional roles in awake behaving animals. Similar to the division of LTMRs into subtypes, there may also be physiologically distinct subtypes of PSDCs, given their various contributions to the DCN described here and their heterogeneous response properties in cats^5^. Future work will address potential PSDC subdivisions, how features are represented across a broader range of DCN PN subtypes, such as those that project to the cerebellum and higher-order thalamic nuclei, and how PSDCs contribute to touch in different behaviors.

We observed striking somatotopic alignment of the direct and indirect dorsal column pathway inputs to VPL-PNs in the DCN. This alignment enables rich representations in VPL-PNs without compromising spatial detail. The development of this somatotopy likely requires complex coordination between primary sensory neurons, including both LTMRs and HTMRs, spinal cord dorsal horn circuitry, and the DCN. Developmental activity, either spontaneous or mechanically evoked, may play an essential role in organizing this system, as it does in other sensory systems^49, 50^. Moreover, adult PSDCs can exhibit robust receptive field plasticity^51, 52^, raising the possibility that signal propagation via the indirect dorsal column pathway can be modified by sensory experience. Thus, subdivisions of the dorsal column pathway may not only expand the capacity for coding tactile features across DCN output pathways, but may also introduce hard-wired and flexible components to the repertoire of mechanosensory representation in the earliest stages of the sensory hierarchy.

## EXTENDED DATA FIGURES

**Extended Data Fig. 1:**
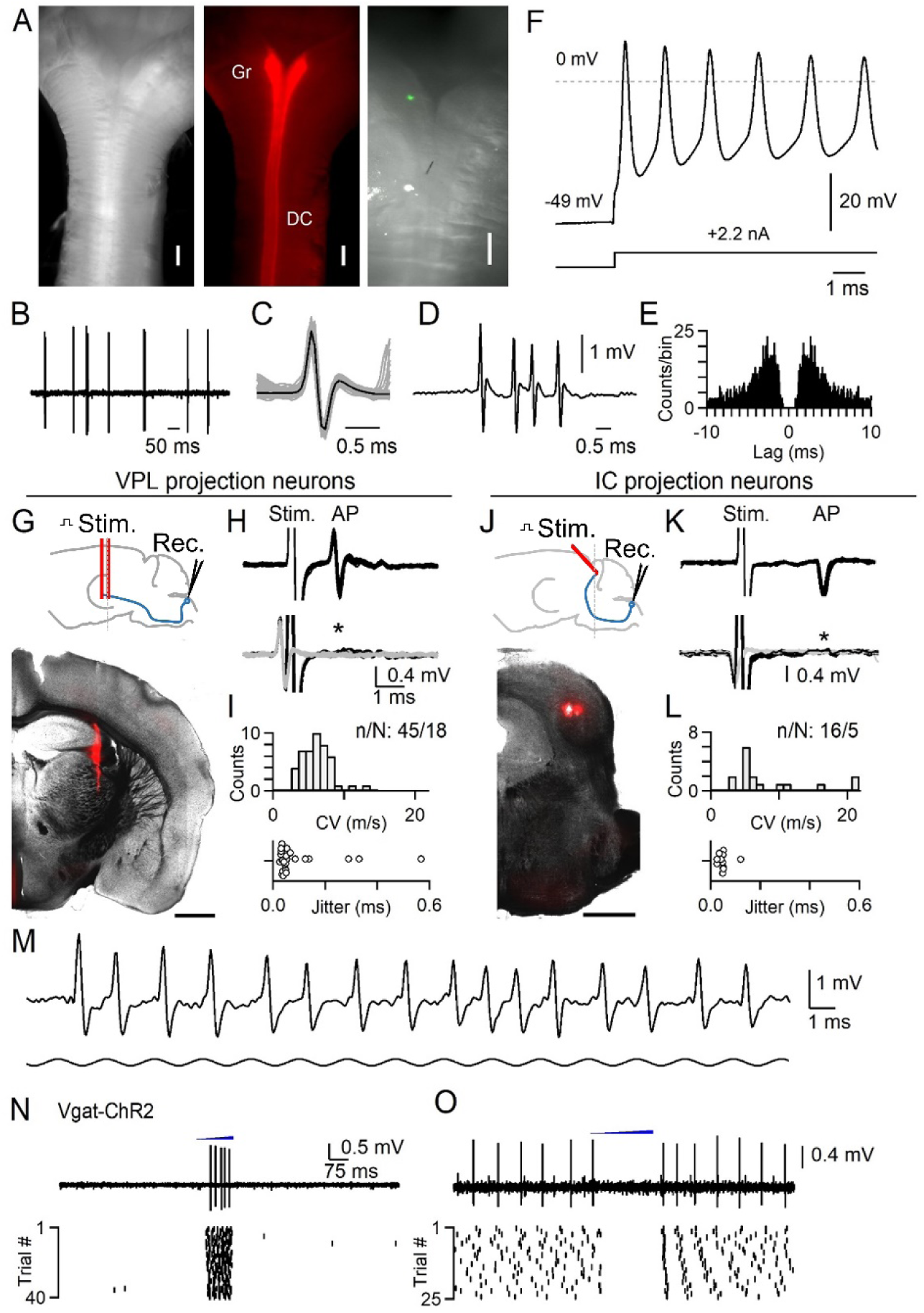
Juxtacellular recording in the DCN, antidromic activation, and optotagging. Juxtacellular recordings were made from units in the gracile nucleus of the DCN to obtain high signal-to-noise recordings of putative single units. Recordings were obtained in urethane-anesthetized mice. Units in the DCN can fire at very high rates up to 1 kHz and can undergo use-dependent changes in action potential waveforms during bursts^54^. Many response properties seen in non-anesthetized preparations are present in anesthetized animals, including a wide distribution of receptive field sizes, entrainment to high-frequency vibration, inhibitory surround receptive fields, and persistent responses to sustained stimuli^46–48^. However, urethane anesthesia will suppress descending input that may modulate the DCN, and may alter polysynaptic circuits that affect DCN activity during different behavioral states. **A.** *<u>Left</u>*: Top view showing the dorsal surface of an isolated brainstem with myelinated dorsal columns forming a Y as they reach the DCN. *<u>Middle</u>*: AAV1-hSyn-Cre was injected into the lumbar cord of a mouse expressing *Rosa26*^LSL-Acr^^1-FRed^, labeling the dorsal columns (DC) and gracile nucleus (Gr). *<u>Right</u>*: Zoomed in view of the gracile nucleus following a recording when the electrode was coated with DiI. The location of the recording site can be seen in green. **B.** Example of spontaneous firing recorded juxtacellularly in the DCN *in vivo*. **C.** Overlaid spikes with single events (gray) and average waveform (black). **D.** Example high frequency burst. **E.** Autocorrelogram for unit in A-D. **F.** Ex vivo intracellular recording of a DCN neuron from an intact brainstem of an adult mouse. Recording was performed in near-physiological conditions (35°C). A current injection was supplied to trigger a brief burst of very high frequency firing. Cells could fire up to 1 kHz bursts with use-dependent changes in action potential waveform as observed *in vivo*. **G.** Identification of VPL-projection neurons (VPL-PNs). *<u>Top</u>*: schematic of preparation, a bipolar electrode was lowered into the VPL. *<u>Bottom</u>*: Example coronal section (100 µm thickness) showing track of the stimulating electrode that was coated with DiI. Scale bar is 1 mm. **H.** *<u>Top</u>*: A unit in the DCN antidromically activated from the VPL. Overlay of 20 trials shows consistent waveform of the stimulus artifact (stim.) and antidromic action potential (AP). The conduction velocity was calculated from the latency to spike following the stimulus artifact with an estimated distance of 9.8 mm. *<u>Bottom</u>*: during a collision test, the stimulus was triggered from the spontaneous firing of the unit. Overlay of 20 trials when electrical stimuli were triggered from spontaneous firing (black). In these trials, stimulation failed to evoke an antidromic spike that would be expected at the latency indicated by the asterisk. Waveform of spontaneous firing in which stimuli were not triggered are overlaid in gray. **I.** *<u>Top</u>*: Histogram of calculated conduction velocities for units antidromically activated from the VPL. Distance from the DCN to VPL was estimated to be 9.8 mm. *<u>Bottom</u>*: Jitter (standard deviation) of latency from units antidromically activated from the VPL. **J.** Identification of IC-projection neurons (IC-PNs). *<u>Top</u>*: schematic of preparation, a bipolar electrode was angled at 45° in order to avoid the sinus above the IC. *<u>Bottom</u>*: Example coronal section (100 µm thickness) showing track of the stimulating electrode that was coated with DiI. Scale bar is 1 mm. **K.** Same as in H, but for a unit antidromically activated from the IC. **L.** Same as in I, but for units activated from the IC. Distance from the DCN to IC was estimated to be 9.4 mm. IC units are those that were classified as vibration-tuned only (see Extended Data Figure 2). **M.** Juxtacellular recording of a single DCN unit *in vivo* showing response to high frequency (500 Hz) vibration of the hindlimb. The unit could entrain to single cycles of the vibration and underwent use-dependent changes in waveform. Vibration begins 6 ms prior to displayed time window. **N.** Optotagging of inhibitory interneurons in the DCN in a Vgat-ChR2 mouse. A fiber optic was positioned above the DCN (400 µm diameter, 1 mm away). Light from a fiber coupled LED was ramped for 200-300 ms (470 nm, 1-2 mW/mm^2^). Optotagged units fired during the ramp. A single trial (*<u>top</u>*) and raster of all trials (*<u>bottom</u>*) for an optotagged unit are shown. **O.** Units that were not optotagged were suppressed by light ramps, or fired rebounds after termination of the ramp.

**Extended Data Fig. 2:**
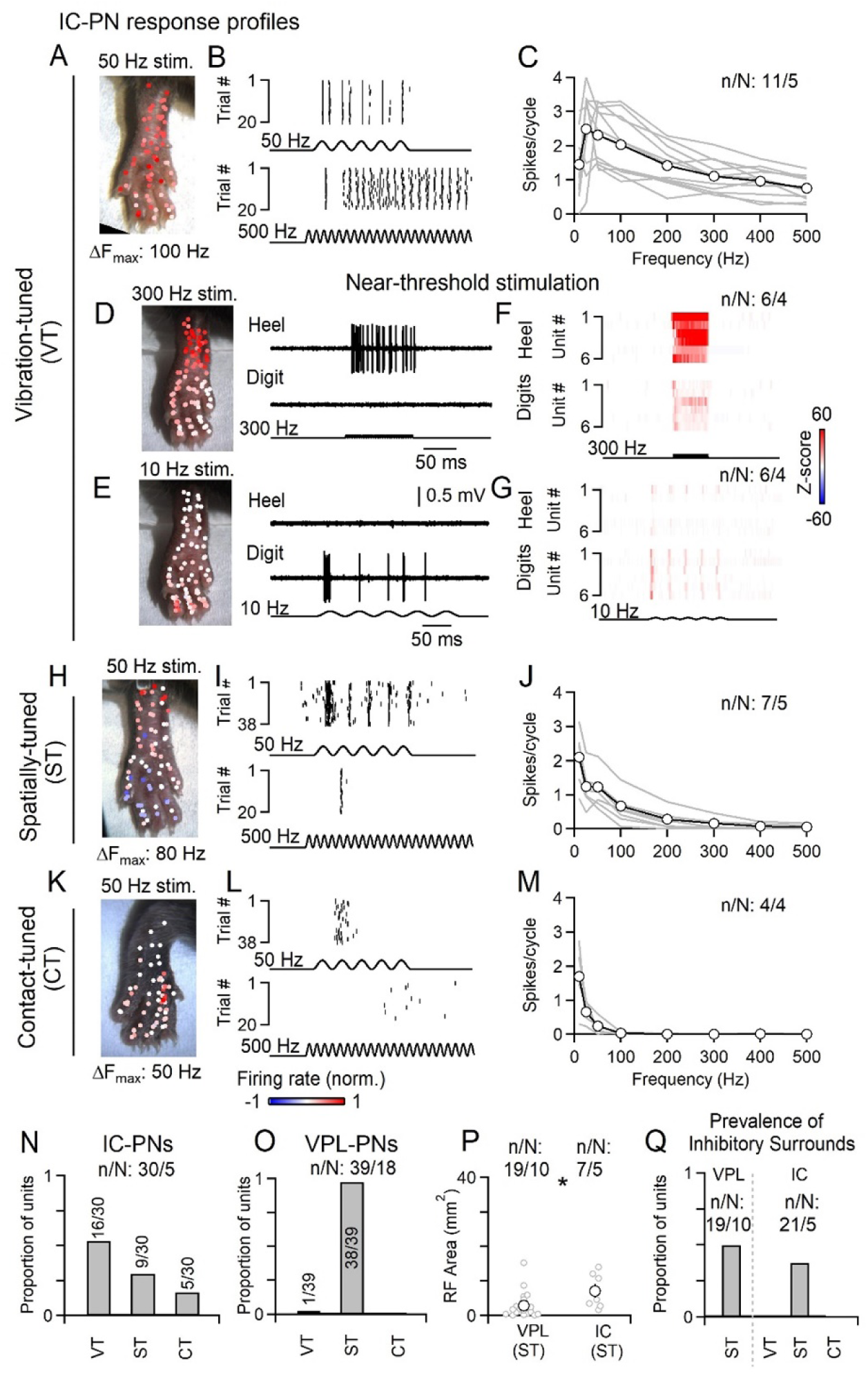
IC projection neuron subtypes. The DCN is composed of several types of projection neurons and local interneurons. VPL-PNs are the most abundant, with an estimated proportion of VPL-PN:IC-PN:Vgat-IN of 2:1:1 based on retrograde tracing and genetic labeling in mice^13^. The proportion of VPL-PNs may be larger^12^. Units that could be antidromically activated from the IC fell into three functional types with distinct properties. Vibration-tuned (VT) units have very large receptive fields that include almost the entire hindlimb, they can entrain their firing to very high frequencies (>300 Hz and often up to 500 Hz), and do not have inhibitory surrounds. The broad receptive field and entrainment to high frequencies suggest that they receive strong input from Pacinian corpuscles. Spatially-tuned (ST) units have small receptive fields, typically consisting of 1-2 digits or pads, or a small portion of the heel. They often have inhibitory surrounds, and can entrain their firing to modest vibration frequencies (up to 100-200 Hz). These units cannot entrain their firing to vibration frequencies >300 Hz, even at high forces. Contact-tuned (CT) units have receptive field sizes similar to spatially-tuned units, but are poorly activated by vibration. They are very sensitive and almost exclusively responsive to the initial contact of a stimulus. **A.** Example receptive field of a vibration-tuned IC-PN. Color scale of normalized firing rate shown at bottom. **B.** Example rasters of a vibration-tuned IC-PN unit in response to 50 (*<u>top</u>*) or 500 Hz (*<u>bottom</u>*) vibration. **C.** Responsiveness of vibration-tuned IC-PN units to different vibration frequencies (10-20 mN). Data is shown for individual units (gray) and average across units (black). **D-G**. Vibration-tuned units had different sensitivities to different frequencies of vibration. When reducing the force of vibrations to near-threshold (<10 mN), 300 Hz vibrations evoked more robust responses in the heel, whereas 10 Hz vibrations typically only evoked responses when stimulating the digits. **D.** Receptive field map of near-threshold 300 Hz vibration for an identified vibration-tuned IC-PN. Example trials from the heel (*<u>top right</u>*) and digits (*<u>top left</u>*) are shown. Color scale of normalized firing rate shown at bottom. **E.** Same as in D, but for 10 Hz near-threshold vibration. Color scale of normalized firing rate shown at bottom. **F.** Average histograms for individual units of near-threshold 300 Hz vibration delivered to the heel (*<u>top</u>*) or digits (*<u>bottom</u>*). Color scale of Z-scores shown at right. **G.** Average histograms for individual units of near-threshold 10 Hz vibration delivered to the heel (top) or digits (bottom). Color scale of Z-scores shown at right. **H-J**. Same as A-C, but for spatially-tuned units. **K-M**. Same as A-C, but for contact-tuned units. **N.** Distribution of functional types for units antidromically activated from the IC. **O.** Functional types were generalized to classify VPL-PNs. Units antidromically activated from the VPL were almost all spatially-tuned. Their receptive fields were small and did not phase-lock to high frequency vibration (>300 Hz), even when increasing the amplitude. These units also typically did not elevate their firing in response to high-frequency vibration. 1/39 units were found to be vibration-tuned. **P.** Receptive field area for spatially-tuned units antidromically activated from the IC or VPL. Distributions are significantly different (p < 0.001, Kolmogorov-Smirnov test). Individual units (gray) and average ± SEM (black). **Q.** Proportion of units with inhibitory surrounds for each functional type.

**Extended Data Fig. 3:**
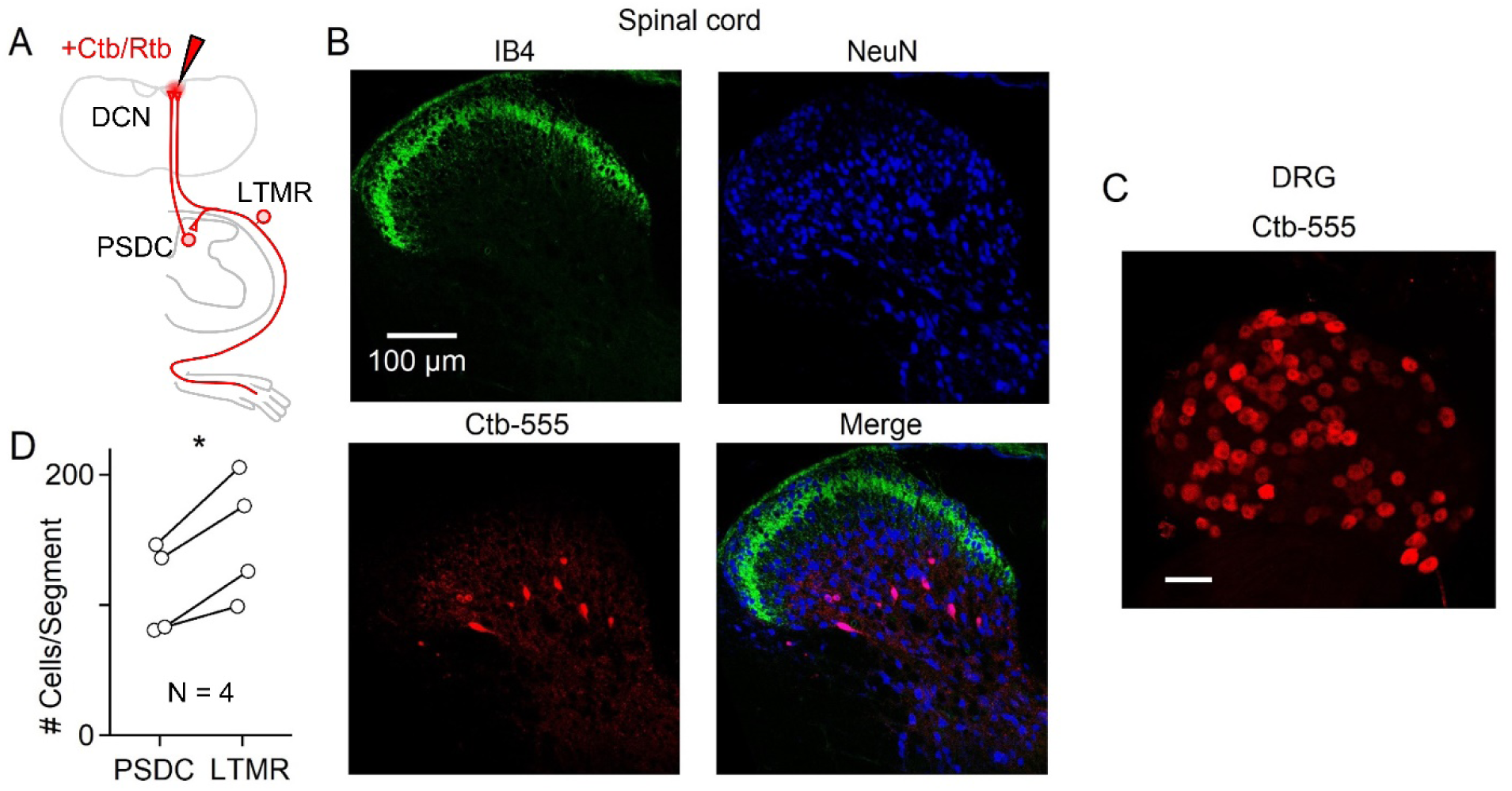
Prevalence and anatomical location of PSDCs. Choleratoxin-B (Ctb) or Retrobeads (Rtb) were injected into the DCN or C1 DC to label PSDCs and Aβ-LTMRs that project to the DCN. Two different tracers were used in order to avoid potential differences in LTMR vs. PSDC uptake efficiency. Small injections were performed in order to avoid off-target labeling in the spinal cord, and to compare the relative number of LTMRs and PSDCs. The number of LTMRs and PSDCs per thoracic spinal segment within the same animal was compared. PSDCs were found in lamina III-IV as described by others^34, 35, 55^. **A.** Schematic showing injection of retrograde tracer into the brainstem and uptake by either PSDCs in the dorsal horn of the spinal cord or Aβ-LTMRs in the DRG. **B.** 10 µm Z-projection of a transverse thoracic spinal cord section (60 µm thick) with immunofluorescence of IB4 (*<u>top left</u>*, green), NeuN (*<u>top right</u>*, blue), Ctb-555 (*<u>bottom left</u>*), and merged images (*<u>bottom right</u>*). **C.** Single plane image of a whole-mount thoracic DRG containing sensory neurons labeled with Ctb-555 from the same animal shown in B. **D.** Comparison of number of Ctb or Rtb labeled cells in the thoracic spinal cord. Each marker pair is the average from one animal, showing the average number of cells labeled in 1-2 DRGs, and the number of PSDCs estimated to exist in one spinal segment (1000 µm length of spinal cord) based on the average number of PSDCs labeled in 10-20 spinal sections (60 µm thickness). P < 0.05, paired t-test.

**Extended Data Fig. 4:**
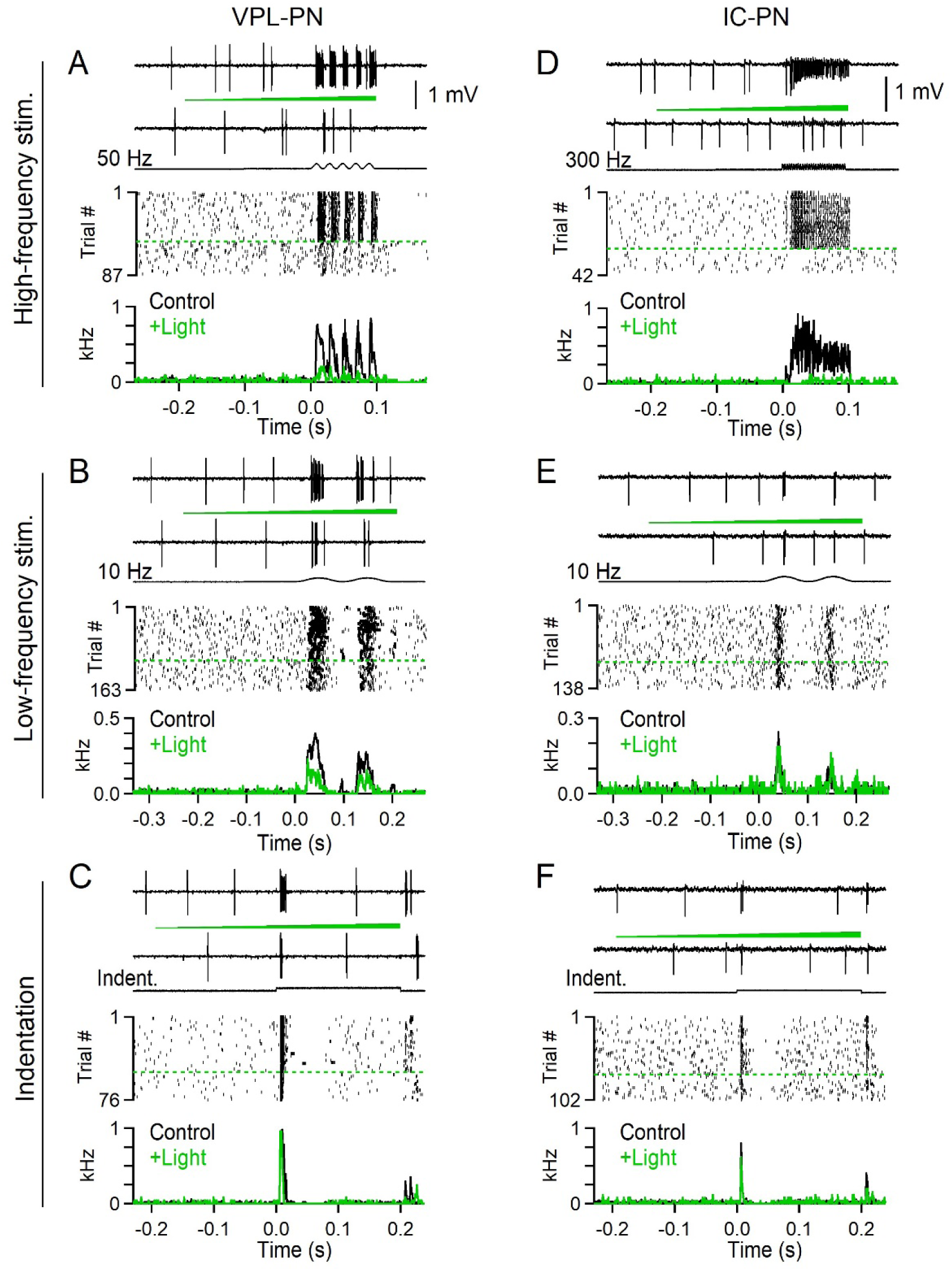
Effects of silencing sensory neuron terminals in the DCN using Acr1 on the response properties of identified projection neurons. Juxtacellular recording of a VPL-PN (left column) and a vibration-tuned IC-PN (right column) in *Avil^Cre^; Rosa26^LSL-Acr^*^1^ animals. In this experiment direct LTMR input is silenced. Mechanical stimuli were applied to the center of the receptive field. On some trials, light (550 nm, 0.46 mW/mm^2^) was applied to the surface of the brainstem using a fiber optic (400 µm diameter, 1 mm away). The light was ramped over the course of 300-400 ms, and mechanical stimuli were delivered in the last 100-200 ms of light. Trials were interleaved but are sorted here for presentation. **A-C**. A DCN unit antidromically activated from the VPL. This unit did not fire in response to high-frequency (≥300 Hz) vibration, even at high forces. **A.** *<u>Top</u>*: Traces of single control and silencing trials. A 100 ms, 50 Hz, 10 mN vibration was applied. Timing of light delivery is denoted by a green ramp. *<u>Middle</u>*: Raster of trials for 50 Hz vibration. Control and light trials are separated by a dashed green line. *<u>Bottom</u>*: Average histogram for this unit when light was off (Control) or on (+Light). Time 0 is the onset of vibration. Bins are 1 ms. **B.** Same unit as in A, but for a 200 ms 10 Hz 10 mN vibration. **C.** Same unit as in A and B, but for a 200 ms 10 mN indentation. **D-F**. A DCN unit that was antidromically activated from the IC. This unit entrained to 300 Hz vibratory stimuli and had a large receptive field. **D.** *<u>Top</u>*: Traces of single control and silencing trials. A 100 ms, 300 Hz, 10 mN vibration was applied. *<u>Middle</u>*: Raster of trials for 300 Hz vibration. Control and light trials are separated by a dashed green line. *<u>Bottom</u>*: Average histogram for this unit when light was off (Control) or on (+Light). Time 0 is the onset of vibration. Bins are 1 ms. **E.** Same unit as in D, but for a 200 ms 10 Hz 10 mN vibration. **F.** Same unit as in D and E, but for a 200 ms 10 mN indentation.

**Extended Data Fig. 5:**
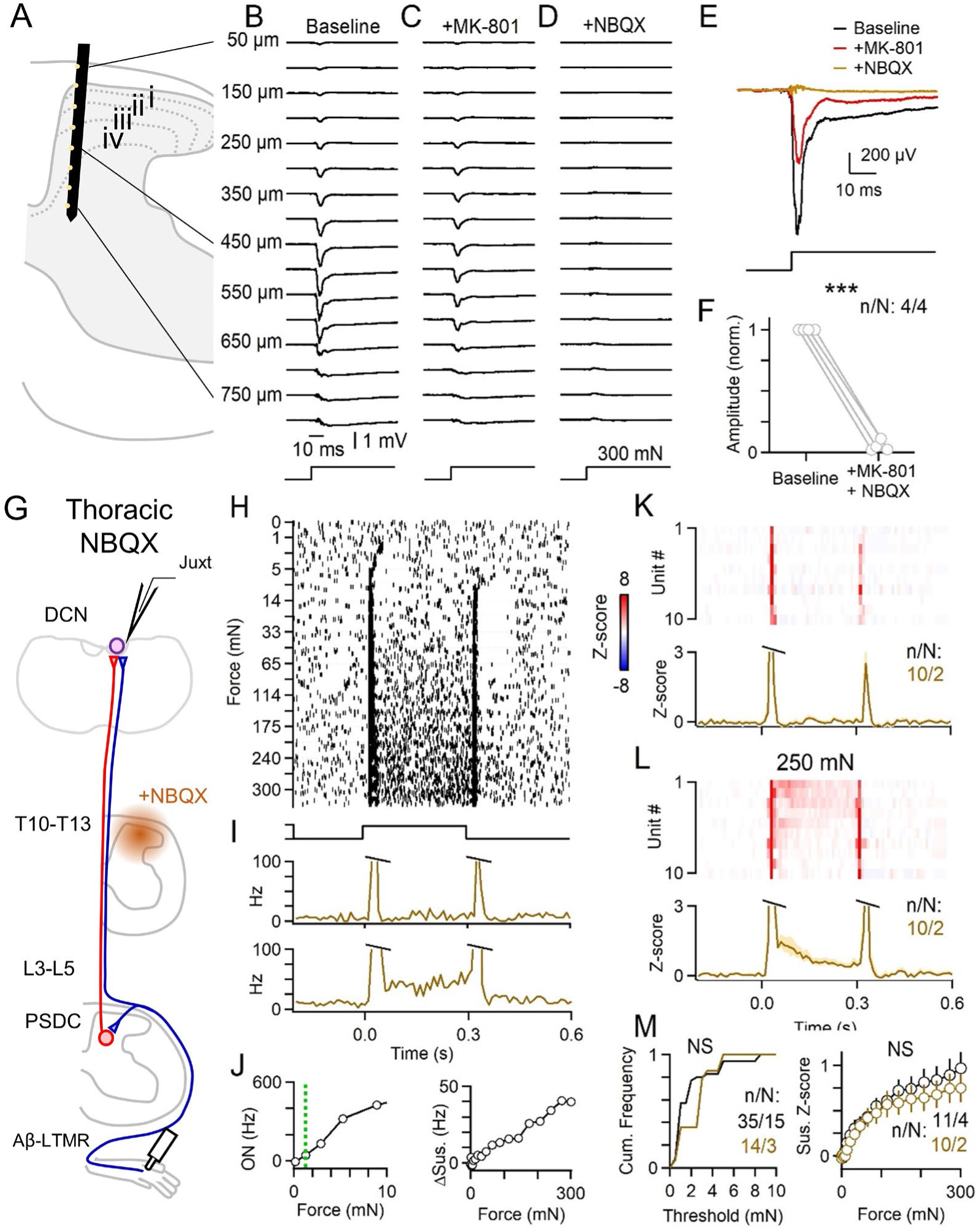
Blockade of glutamatergic transmission suppresses evoked responses locally within the spinal cord. **A-F**. A multi-electrode array (MEA) was inserted into L4 spinal dorsal horn, with recording sites facing medially. Field potentials were recorded in response to a 300 mN step applied to the center of the receptive field on the hindlimb. Following baseline trials, MK-801 (10 mM, 10 µL) was applied to the surface of the spinal cord for 3-5 minutes to block NMDA receptors. The surface was then irrigated with saline and NBQX (10 mM, 20 µL) was applied to block AMPA receptors. **A.** Schematic of approximate recording location. MEA has 16x2 recording sites spaced 50 µm apart. Only a subset of sites are schematized. **B.** Field potentials generated at various depths of the spinal cord in response to a 300 mN step indentation under baseline conditions. **C.** Same recording as in B, but after application of MK-801 to the surface of the spinal cord. **D.** Same recording as in B and C, but after application of NBQX to the surface of the spinal cord. **E.** Overlaid field potentials from a 450 µm depth recording site in response to 300 mN indentation, from recording shown in B-D. Responses following NBQX remained suppressed for the remainder of the recording (30 minutes after NBQX application). **F.** Summary of change in evoked amplitude. Each marker pair is one animal. (Paired t-test, p < 0.01). **G-M.** Pharmacological manipulations can have non-specific effects, as drugs may spread from the site of application or enter the body systemically. We assessed these possibilities by applying glutamatergic antagonists to the thoracic cord (T10-T13) to determine whether hindlimb responses in the DCN may be altered through diffusion of the drug to the DCN or non-targeted regions of the spinal cord. A laminectomy was performed over the T10-T13 spinal segments, and the dura was removed. MK-801 (10 mM, 10 µL) was first applied to the surface of the spinal cord for 5 minutes. The cord was then abundantly irrigated with saline. NBQX (10 mM, 20 µL) was then applied to the surface and gelfoam was placed on top and kept moist with saline. **G.** Schematic of experimental setup. **H.** Raster of example DCN unit in response to step indentations of various forces (0-300 mN) in the presence of NBQX applied to T10-T13 spinal segments. **I.** Histogram of average responses to 10 mN (*<u>top</u>*) and 250 mN (*<u>bottom</u>*) for unit shown in H-I. **J.** *<u>Left</u>*: Max on response (0-20 ms) vs. indentation force. Dashed green line indicates threshold. *<u>Right</u>*: Average sustained firing (50-250 ms) vs. force. **K.** *<u>Top</u>*: Response of DCN units to 10 mN step indentation in the presence of NBQX. *<u>Bottom</u>*: Average across DCN units to 10 mN indentation in the presence of NBQX. **L.** Same as K, but for 250 mN step indentation. **M.** *<u>Left</u>*: Mechanical threshold of on-response for units in the presence (brown) or absence (black) of NBQX over T10-T13 spinal segments. Distributions are not significantly different (Kolmogorov-Smirnov test, p = 0.25). *<u>Right</u>*: Average ± SEM of sustained firing rate as a function of indentation force across cells in the presence (brown) or absence (black) of NBQX over T10-T13. (Kolmogorov-Smirnov test, p = 0.76).

**Extended Data Fig. 6:**
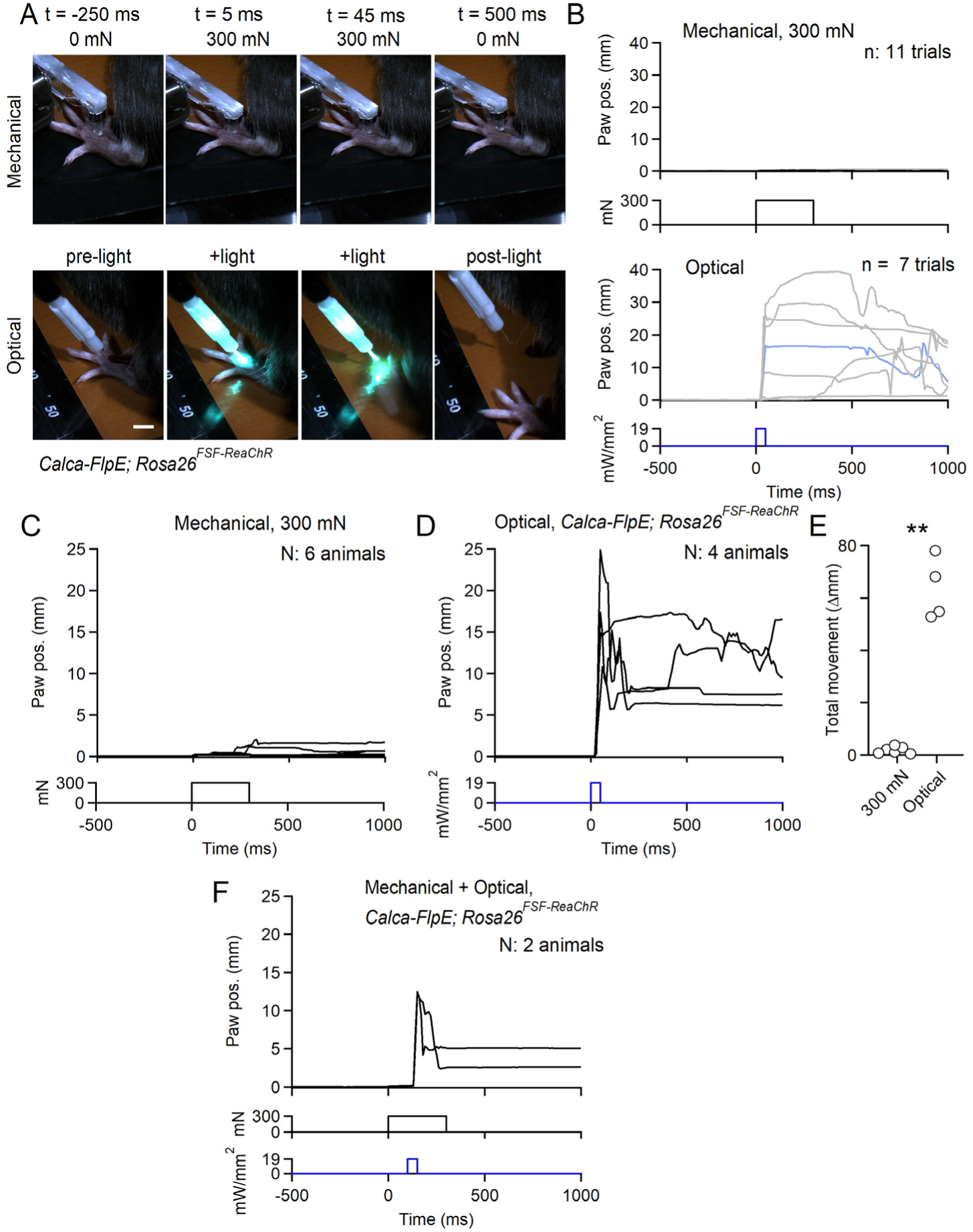
Mechanical stimuli used for physiology experiments do not evoke nocifensive responses. Indentation at high forces (300 mN) using a large, 1 mm probe tip was not perceived as painful when applied to the finger tip of human experimenters (J.T. and D.D.G.). We tested whether this same high force stimulus was noxious for mice. Awake C57Bl/J6 or *Calca*-*FlpE*; *Rosa26^FSF-ReaChR^* animals were head-fixed and otherwise unrestrained. Stimuli were delivered to the paw and the mouse was free to withdraw its paw or attempt to avoid the stimulus. Animals were allowed to habituate for 5-10 minutes. Weak (10-20 mN) indentations were first delivered to the hindpaw to acclimate the animal to the 1 mm probe. Then the amplitude was elevated to 300 mN. The first indentation trial was discarded to avoid capturing startle. 4-15 trials were then collected to determine whether the mouse withdrew its paw. In *Calca*-*FlpE*; *Rosa26^FSF-ReaChR^* animals, the series of indentation trials were then followed by a series of optical stimulation trials. A 400 µm fiber delivered a ∼20 mW/mm^2^ 50 ms pulse of light to the dorsal hindpaw. The first trial was discarded, and 3-7 trials were then collected. See also Supplemental Video 1. **A.** Video frames acquired during either indentation (top) or optical activation of *Calca*^+^ afferents (bottom) in the same mouse. Scale bar is 5 mm. **B.** Plot of paw displacement relative to baseline for 300 mN 300 ms indentation for the mouse shown in A (top), and plot of paw displacement relative to baseline for 50 ms optical stimulation for the same experiment shown in A (bottom). Highlighted blue trace is from trial shown in A. **C.** Average paw displacements (paw position) for 6 mice in response to 300 mN indentation. **D.** Average paw displacements (paw position) for 4 mice in response to optical stimulation of *Calca*^+^ afferents. **E.** Total average movement (distance paw traveled) during mechanical or optical stimulation. Each marker is the average of one animal. Kolmogorov-Smirnov test, p < 0.01. **F.** Lack of paw withdrawal following 300 mN mechanical stimulation was not due to constraint of the paw by the indenter. In two *Calc*a-*FlpE*; *Rosa26^FSF-ReaChR^* animals, a 300 mN indentation was delivered. 100 ms after the onset of indentation, light was also delivered to the paw. Optical stimulation triggered paw withdrawal in the presence of a 300 mN force pressing down on the paw.

**Extended Data Fig. 7:**
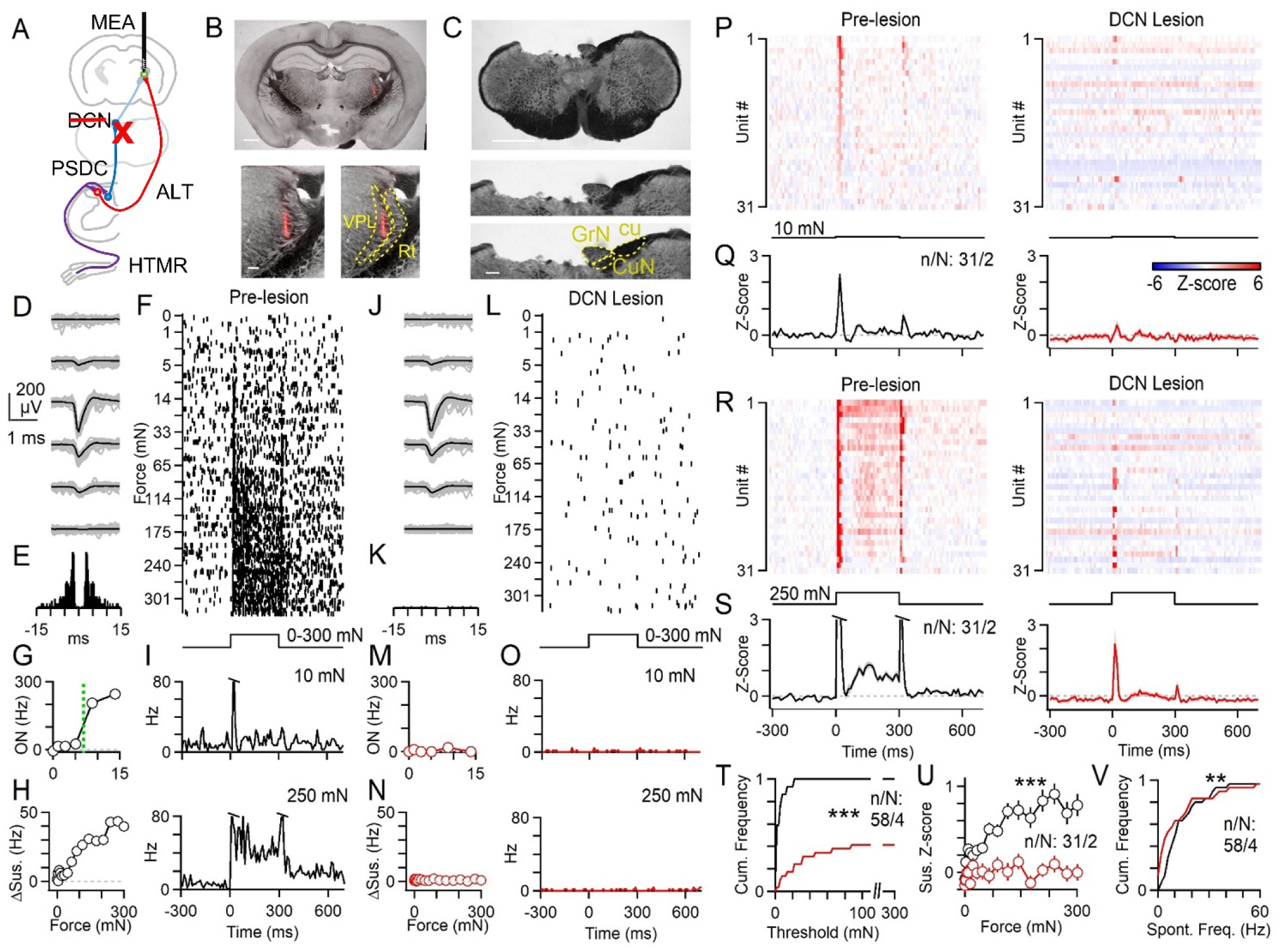
The DCN supplies high force information to the middle VPL (mVPL). The primary somatosensory thalamus can be subdivided into rostral, middle and caudal regions^56^. Rostral regions may receive more proprioceptive input and have large receptive fields. The middle region has cutaneous responses with detailed spatial receptive fields. The caudal region has diffuse receptive fields and may be more responsive to intense nociceptive stimuli. We asked whether the mVPL can encode the intensity of sustained stimuli, and whether such coding arises from the anterolateral tract (ALT) or the dorsal column system. We recorded from the mVPL in urethane-anesthetized mice and lesioned the DCN to measure how mechanically-evoked responses change. The dorsal column system crosses just rostral to the DCN, whereas the ALT crosses within the spinal cord and runs laterally. Other spinothalamic and ascending neurons also cross within the spinal cord and run along the ventral aspect of the brainstem, and should be spared by lesioning the DCN. However, a potential caveat is that lesioning the DCN may disrupt the corticospinal tract, which passes just ventrally to the DCN toward the spinal cord. **A.** Schematized experimental configuration. A 32-channel multi-electrode array (MEA) was inserted into the middle VPL of the right thalamus. Mechanical indentation of the contralateral (left) hindpaw was performed before and after lesion of the left DCN by aspiration. The lesion should disrupt DCN input from reaching the VPL, but spare input from ALT neurons. **B.** Coronal section of the brain (50 µm) following recording with DiI labeling from the MEA in the right VPL (top). Scale bar is 1 mm. Inset shows magnified view of the thalamus (bottom left) and inferred location of VPL and thalamic reticular nucleus (Rt, bottom right). Scale bar is 1 mm (top) and 100 µm (bottom) **C.** Coronal section of the brainstem showing the location of the lesion. The dorsal brainstem was aspirated at the level of the caudal DCN on the left side (top). Inset shows magnified view of the lesion site, and location of the gracile nucleus (GrN) that relays information from the hindpaw, and also the cuneate tract (cu) and some of the cuneate nucleus (CuN). Scale bar is 1 mm (top) and 200 µm (bottom) **D.** Waveform for a unit recorded across four channels of the MEA prior to the DCN lesion with a subset of single events (gray) and overlaid average (black). **E.** Autocorrelogram for unit shown in D-O prior to the lesion. **F.** Raster of firing for unit shown in D-O in response to a 300 ms step indentation of varying force. Trials were interleaved, but are shown sorted for presentation. **G.** Maximum firing rate for the onset of step indentation (0-50 ms) as a function of indentation force for unit shown in D-O prior to lesion. Dashed green line indicates the measured threshold. **H.** Average firing rate during the sustained component of the step indentation (150-300 ms) for unit shown in D-O prior to lesion. **I.** Time course of firing rate for unit shown in D-O in response to indentation for 10 mN (top) and 250 mN (bottom) prior to lesion. **J-O**. Same as D-I, but for the same unit after DCN lesion. **P.** Average histograms for all mVPL units in response to 10 mN 300 ms indentation prior to lesion (left) and for the same units after DCN lesion (right). Units are sorted by strength of on-response to 10 mN indentation. **Q.** Average of all mVPL units in response to 10 mN indentation before (left) and after (right) lesion of the DCN. **R.** Average histograms for all mVPL units in response to 250 mN 300 ms indentation prior to lesion (left) and for the same units after DCN lesion (right). Units are same as shown in P, but sorted by strength of sustained response at 250 mN. **S.** Average of all mVPL units in response to 250 mN indentation before (left) and after (right) lesion of the DCN. **T.** Cumulative histogram of on-response threshold for the same units before (black) and after (red) lesion of the DCN. About 60% of units could no longer be activated by indentations up to 300 mN following DCN lesion. Kolmogorov-Smirnov test, p < 0.01. **U.** Average ± SEM of sustained response of all units as a function of indentation force before (black) and after (red) lesion of the DCN. Kolmogorov-Smirnov test, p < 0.01. **V.** Cumulative histogram of spontaneous firing rates for same units before (black) and after (red) lesion of the DCN. Kolmogorov-Smirnov test, p < 0.01.

**Extended Data Fig. 8:**
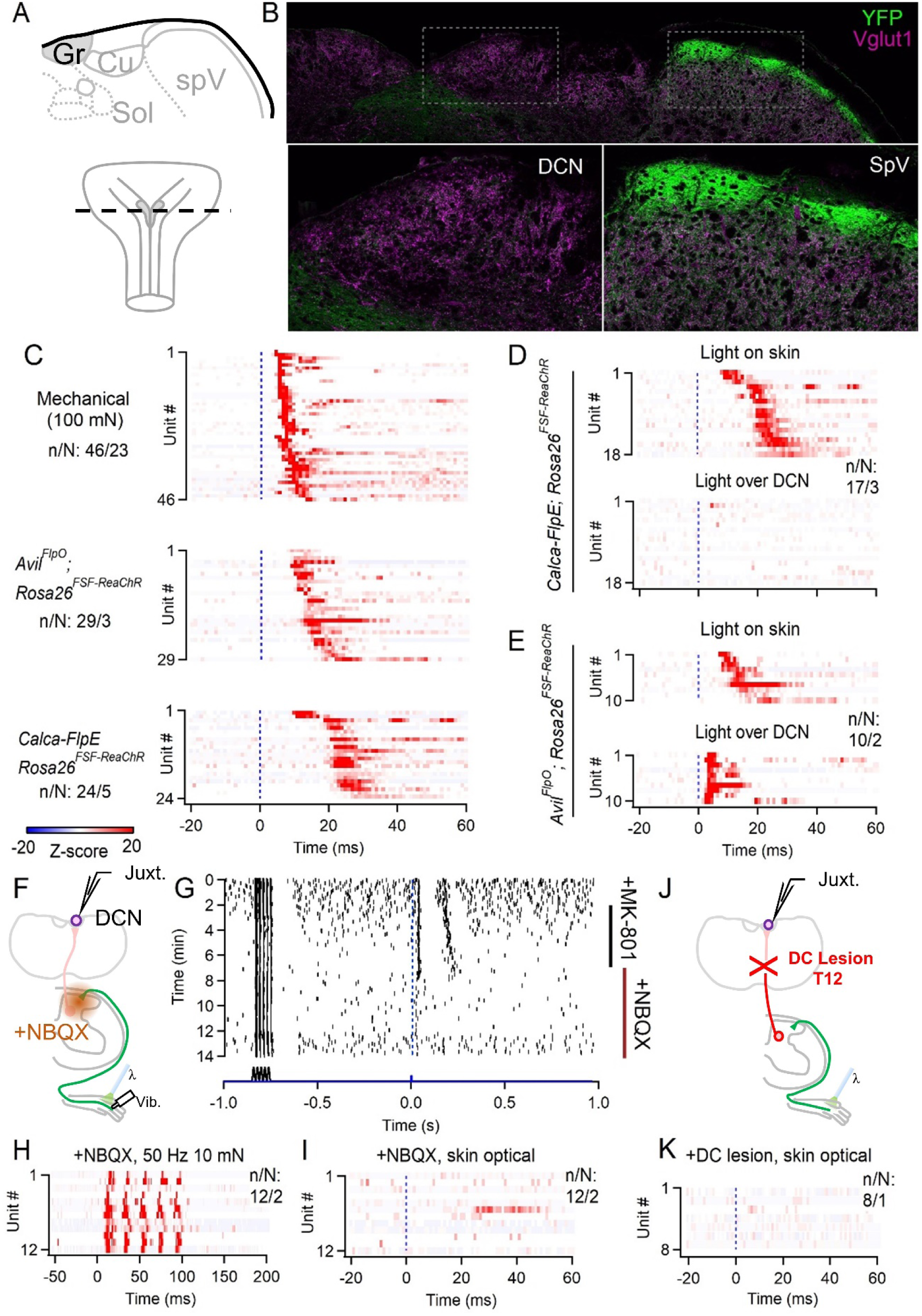
*Calca*^+^ HTMRs do not project directly to the DCN but activation of their endings in the skin drives long latency responses. *Calca* is expressed in a broad set of sensory neurons. These include mechanically sensitive, polymodal receptors, and thermoreceptors that have A or C-fiber conduction velocity; *Calca*^+^ HTMRs have higher mechanical thresholds than Aβ-LTMRs^40–43^. *Calca*^+^ DRG neurons typically do not project to the DCN, although previous studies have shown ∼10% of DRG neurons retrogradely labeled from the DCN express *Calca*^37^. Others have observed sparse *Calca*^+^ fibers within the DCN^44, 45^. **A.** Schematic representation of a coronal section of brainstem containing the gracile and cuneate nucleus of the DCN (Gr, Cu), and the neighboring solitary nuclei (Sol) and spinal trigeminal nucleus (spV). The rostro-caudal area of the brainstem is shown at bottom. **B.** Immunofluorescence of Vglut1 (magenta) and ReaChR-YFP (green) in a brainstem section of a *Calca-FlpE; Rosa26^FSF-ReaChR^* animal showing the dorsal brainstem section corresponding to A (*<u>top</u>*). Insets of the gracile (*<u>bottom left</u>*) and spinal trigeminal (*<u>bottom right</u>*) are shown for comparison. YFP^+^ Fibers are densely labeled in the dorsal spinal trigeminal, but very few were detected in the gracile. **C.** DCN neurons were juxtacellularly recorded in the gracile and the latencies of responses to various stimuli were measured. DCN units had short latency responses to strong mechanical indentation beginning at time 0, in wildtype animals (*<u>top</u>*). In *Avil^FlpO^*; *Rosa26^FSF-ReaChR^* randomly recorded DCN units also had short latency responses to optical activation of sensory neurons in the skin (*<u>middle</u>*). In *Calca-FlpE; Rosa26^FSF-ReaChR^* animals optical activation of *Calca*^+^ endings in the skin evoked longer latency responses in randomly recorded DCN units (*<u>bottom</u>*). **D.** Random DCN units and their average response to light activation of *Calca*^+^ endings in the skin in *Calca-FlpE*; *Rosa26^FSF-ReaChR^* animals (*<u>top</u>*). In the same units, light was also delivered directly over the DCN (*<u>bottom</u>*). Light over the DCN evoked weak responses in 2/19 units. **E.** Random DCN units for *Avil^FlpO^*; *Rosa26^FSF^*^-*ReaChR*^ animals in which all sensory neurons express ReaChR. Robust firing could be evoked in all recorded DCN units when light was delivered over the DCN (*<u>bottom</u>*). **F.** Random DCN neurons were recorded and light was delivered to the skin in *Calca-FlpE*; *Rosa26^FSF-ReaChR^* animals. The same spot on the skin was also stimulated using a gentle (10-20 mN) vibration (50 Hz, 100 ms). Synaptic blockers were then applied to the lumbar spinal cord to measure their effect on optically- and mechanically-evoked responses. **G.** Raster of a random DCN unit and its response to 50 Hz vibration and optical activation of *Calca*^+^ endings in the skin. After a baseline period, the NMDAR antagonist MK-801 was applied to the lumbar spinal cord (10 mM, 10 µL). After 5 minutes, the MK-801 was washed away with saline, and NBQX was added to the spinal cord (10 mM, 10 µL). **H.** Average histograms of vibration responses of random DCN units in which synaptic blockers were applied to the spinal cord. **I.** Same units shown in H and their response to optical activation of *Calca*^+^ endings in the skin following application of synaptic blockers to the spinal cord. **J.** Random units in the DCN were recorded in a *Calca-FlpE*; *Rosa26^FSF-ReaChR^* animal. The area of the gracile responding to the hindlimb was located and the DC at T12 was lesioned. Responses were then measured to optical activation of *Calca*^+^ endings in the skin. **K.** Average responses in random DCN units in hindlimb region of the gracile to optical activation of *Calca*^+^ endings in the skin after lesioning the DC at T12. The entire hindpaw was sampled for each unit to ensure the receptive field was not missed.

**Extended Data Fig. 9:**
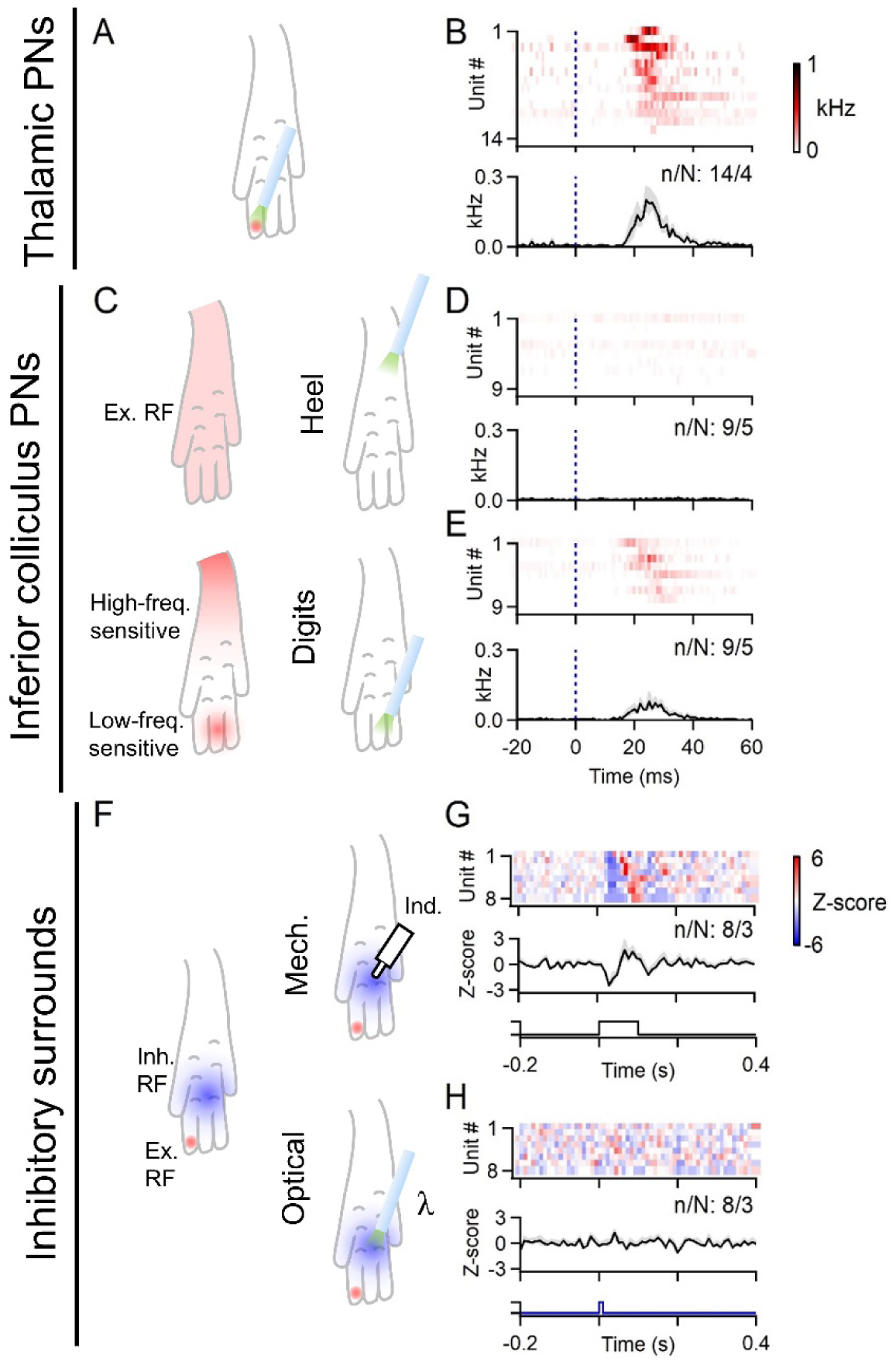
*Calca*^+^ HTMRs can drive IC-PNs, but not inhibitory surrounds. All experiments were performed in *Calca*-*FlpE*; *Rosa26^FSF-ReaChR^* animals. **A.** Schematic representation of a typical VPL-PN excitatory receptive field and location of light stimulus delivered. **B.** Average firing rate of identified VPL-PNs in response to optical activation of *Calca*^+^ endings in the skin (*<u>top</u>*). Average evoked firing rate across all VPL neurons following optical stimulation of *Calca*^+^ cutaneous endings (*<u>bottom</u>*). Bins are 1 ms. Color scale for firing rate shown at right. **C.** The receptive fields of vibration-tuned IC-PNs typically span the entire paw (*<u>top left</u>*). However, with very low stimulus intensities, it was revealed that the heel is more sensitive to high-frequency vibration, whereas the toes are more sensitive to low frequency vibration (*<u>bottom</u> <u>left</u>*; see Extended Data Fig. 2). Light was thus delivered to either the heel (*<u>top right</u>*), or the most sensitive area of the digits (*<u>bottom right</u>*). **D.** Response of individual vibration-tuned IC-PNs in response to optical stimulation of *Calca*^+^ afferents in the heel (*<u>top</u>*), and average response across units (*<u>bottom</u>*). Bins are 1 ms. Color scale for firing rate shown in B. **E.** Response of individual vibration-tuned IC-PNs in response to optical stimulation of the digits (*<u>top</u>*), and average response across units (*<u>bottom</u>*). Units are the same as those shown in D. **F.** The inhibitory surround of a DCN unit was stimulated mechanically using 10-20 mN indentation (*<u>top right</u>*). The same part of the inhibitory surround was then also stimulated optically (*<u>bottom right</u>*). **G.** Average responses of individual DCN units to 10-20 mN indentation delivered to their inhibitory surrounds (*<u>top</u>*), and average inhibitory response across all units (*<u>bottom</u>*). Bins are 10 ms. Color scale Z-score shown at right. **H.** Average responses of individual DCN units to optical pulse delivered to their inhibitory surrounds (*<u>top</u>*), and average response across all units (*<u>bottom</u>*). Units are the same as those shown in G.

**Extended Data Fig. 10:**
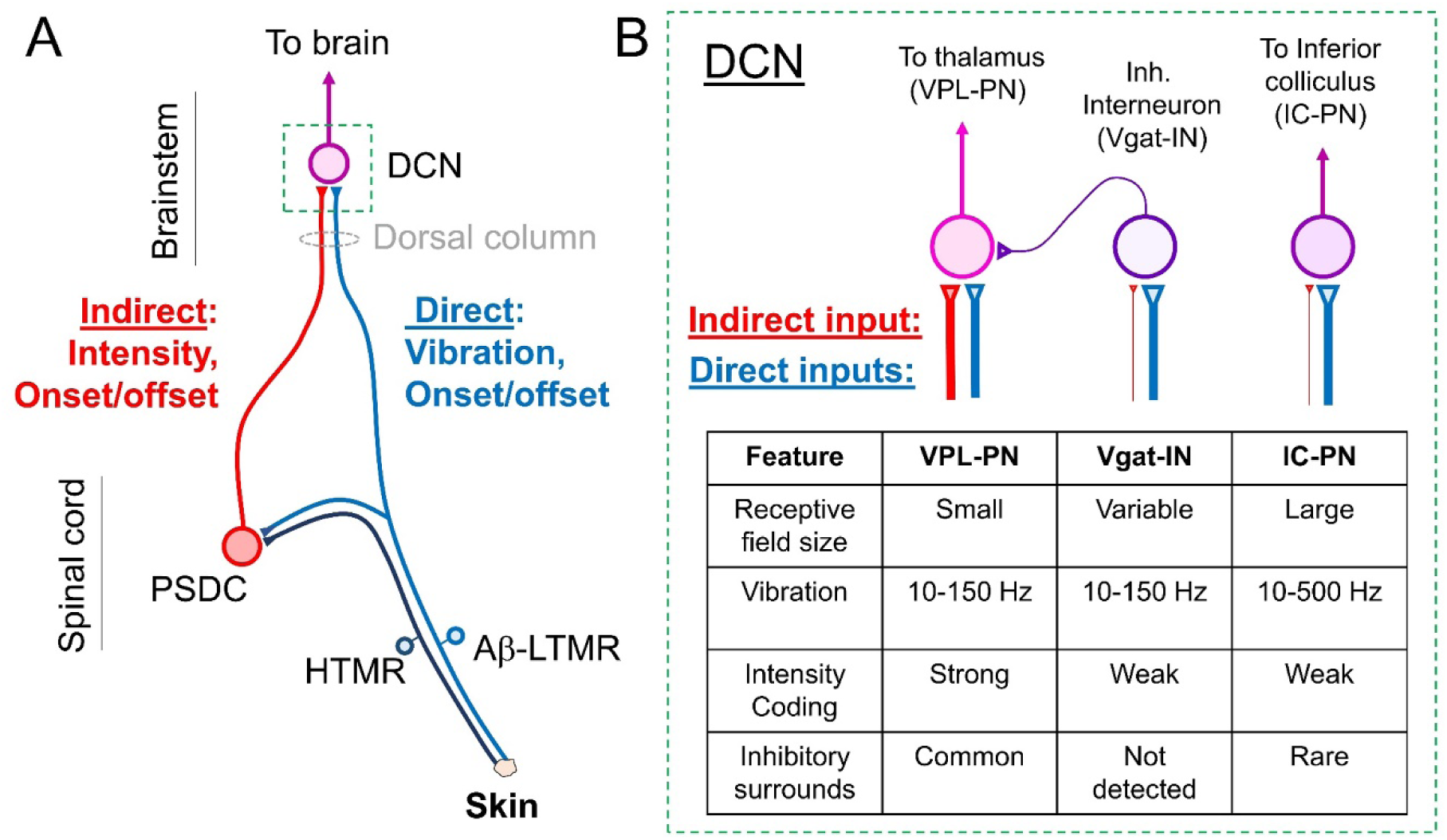
A model for sensory representation in the hindlimb dorsal column system. **A.** Mechanical stimuli in a small area of skin are detected by LTMRs and HTMRs. LTMRs ascend the dorsal column and synapse onto neurons in the DCN, forming the direct pathway. LTMRs also form collaterals within the spinal cord dorsal horn. PSDCs in the spinal cord dorsal horn receive mechanical information from LTMRs and HTMRs, either directly or through local interneurons. PSDC axons then ascend the dorsal column, forming the indirect pathway. PSDCs of the indirect pathway signal the intensity of sustained stimuli and the onset/offset of stimuli, as well as low-frequency vibratory stimuli. LTMRs of the direct pathway signal rapid movements and high-frequency vibratory stimuli, in addition to the onset/offset of stimulation. These two pathways converge somatotopically such that individual DCN neurons can encode multiple features in a small patch of skin. **B.** Different contributions from PSDCs and LTMRs drive distinct response properties in DCN neuron types. *<u>Left</u>*: VPL-PNs have small RF sizes and commonly have inhibitory surrounds driven by DCN Vgat-INs. Direct LTMR input drives phase-locked spiking up to ∼150 Hz. VPL- PNs can also commonly encode sustained stimuli at high forces mediated by indirect PSDC input. Thus, VPL-PNs receive prominent direct and indirect input. *<u>Middle</u>:* Local Vgat-INs have varying receptive field sizes and can entrain to vibrations up to 150 Hz. They can, in some cases, be activated by high-frequency stimulation, but typically do not entrain their firing. Vgat-INs rarely fire in response to the sustained component of stimuli and thus poorly encode the intensity of sustained stimuli. Inhibitory surrounds in other DCN neurons that are generated by Vgat-INs are not strongly driven by indirect input, and are not driven by *Calca*^+^ HTMR activation. *<u>Right</u>:* The most common type of IC-PNs (Vibration-tuned) have very large receptive field sizes and can entrain to high-frequency vibratory stimuli up to 500 Hz. These high-frequency responses are likely generated exclusively by direct LTMR (Aβ-RA2-LTMR) input. They can encode sustained stimuli, but not as well as VPL-PNs. They receive some input from HTMRs, likely through PSDCs, but only to a subset of their receptive field.

**Extended Data Table 1:** number of experiments

| Figure | Condition/Experiment | n (units) | N (animals) |
| --- | --- | --- | --- |
| 1D | VPL-PN frequencies | 6 | 4 |
| 1E | VPL-PN indentation | 22 | 10 |
| 1I | IC-PN frequencies | 11 | 5 |
| 1J | IC-PN indentation | 10 | 5 |
| 1N | Vgat-IN frequencies | 10 | 3 |
| 1O | Vgat-IN indentation | 12 | 3 |
| 1P | VPL-PN RF area | 19 | 10 |
| 1P | IC-PN RF area | 11 | 4 |
| 1P | Vgat-IN RF area | 12 | 3 |
| 1Q | VPL-PN inh. Surr. | 19 | 10 |
| 1Q | IC-PN inh. Surr. | 11 | 4 |
| 1Q | Vgat-IN inh. Surr. | 12 | 3 |
| 1R | VPL-PN Spont. Rate | 23 | 10 |
| 1R | IC-PN Spont. Rate | 12 | 5 |
| 1R | Vgat-IN Spont. Rate | 12 | 3 |
| 2C | 10 Hz | 7 | 2 |
| 2C | 50 Hz | 7 | 3 |
| 2C | 300 Hz | 5 | 1 |
| 2D-E | Indentation | 10 | 1 |
| 2H | 10 Hz | 14 | 4 |
| 2H | 50 Hz | 14 | 4 |
| 2H | 300 Hz | 7 | 2 |
| 2I-J | Indentation | 10 | 3 |
| 2L-N | Inh. Surr. 10 Hz | 8 | 3 |
| 2L-N | Inh. Surr. 50 Hz | 10 | 6 |
| 2L-N | Inh. Surr. Indent. | 11 | 3 |
| 3C | 10 Hz Control | 8 | 2 |
| 3C | 50 Hz Control | 14 | 4 |
| 3C | 300 Hz Control | 8 | 2 |
| 3C | 10 Hz NBQX | 5 | 1 |
| 3C | 50 Hz NBQX | 11 | 2 |
| 3C | 300 Hz NBQX | 5 | 1 |
| 3J-K | Indentation Ctrl. | 11 | 4 |
| 3J-K | Indentation NBQX (L3-L5) | 10 | 4 |
| 3L | Threshold Ctrl. | 35 | 15 |
| 3L | Threshold NBQX (L3-L5) | 19 | 7 |
| 3M | Sustained firing Ctrl. | 11 | 4 |
| 3M | Sustained firing NBQX (L3-L5) | 10 | 4 |
| 4C,E | VPL-PN 100 mN indent. | 23 | 9 |
| 4C,E | VPL-PN Calca input | 14 | 4 |
| 4K,L | Mech./Optical RFs | 7 | 2 |
| ED1 I | VPL-PN CV, jitter | 45 | 18 |
| ED1 L | IC-PN CV, jitter | 16 | 5 |
| ED2 C | IC-PN VT frequencies | 11 | 5 |
| ED2 F,G | IC-PN heel vs. digit | 6 | 4 |
| ED2 J | IC-PN ST frequencies | 7 | 5 |
| ED2 M | IC-PN CT frequencies | 4 | 4 |
| ED2 N | IC-PN VT | 16 | 5 |
| ED2 N | IC-PN ST | 9 | 5 |
| ED2 N | IC-PN CT | 5 | 3 |
| ED2 O | VPL-PN VT | 1 | 1 |
| ED2 O | VPL-PN ST | 38 | 17 |
| ED2 O | VPL-PN CT | 0 | 0 |
| ED2 P | VPL-PN ST RF area | 19 | 10 |
| ED2 P | IC-PN ST RF area | 7 | 5 |
| ED2 Q | IC-PN VT inh. Surr. | 11 | 4 |
| ED2 Q | IC-PN ST inh. Surr. | 7 | 5 |
| ED2 Q | IC-PN CT inh. Surr. | 3 | 3 |
| ED2 Q | VPL-PN ST inh. Surr. | 19 | 10 |
| ED3 D | LTMR/PSDC count | N/A | 4 |
| ED5 F | Field potentials +NBQX | N/A | 4 |
| ED5 K-L | NBQX T10-T13 | 10 | 2 |
| ED5 M | NBQX T10-T13 threshold | 14 | 3 |
| ED5 M | NBQX T10-T13 sustained | 10 | 2 |
| ED6 C, E | Behavior Mechanical | 2 BL6, 4 *Calca-FlpE* | 6 |
| ED6 D, E | Behavior Optical | 4 *Calca-FlpE* | 4 |
| ED6 F | Behavior Mechanical+Optical | 2 *Calca-FlpE* | 2 |
| ED7 P-S | VPL indentation (pre/post) | 31 | 2 |
| ED7 T | VPL threshold (pre/post) | 58 | 4 |
| ED7 U | VPL sustained (pre/post) | 31 | 2 |
| Text | DCN spont. rate (pre/post) | 16 | 2 |
| ED7 V | VPL spont. rate (pre/post) | 58 | 4 |
| ED8 C | DCN 100 mN indent. random | 46 | 23 |
| ED8 C | *Avil<sup>FlpO</sup>; Rosa26<sup>FSF-ReaChR</sup>* | 29 | 3 |
| ED8 C | *Calca-FlpE; Rosa26<sup>FSF-ReaChR</sup>* | 24 | 5 |
| ED8 D | *Calca-FlpE; Rosa26<sup>FSF-ReaChR</sup>* | 18 | 3 |
| ED8 E | *Avil<sup>FlpO</sup>; Rosa26<sup>FSF-ReaChR</sup>* | 10 | 2 |
| ED8 H-I | NBQX; *Calca-FlpE* | 12 | 2 |
| ED8 K | DC lesion; *Calca-FlpE* | 8 | 1 |
| ED9 B | VPL-PN optical | 14 | 4 |
| ED9 D-E | IC-PN optical | 9 | 5 |
| ED9 G-H | Inh. Surrounds optical | 8 | 3 |

**Extended Data Table 2:** statistics

| Figure | Experiment | p-value | Statistical test |
| --- | --- | --- | --- |
| 1P | RF area; VPL/IC-PN | **<0.0001** | Kolmogorov-Smirnov |
| 1P | RF area; VPL-PN/Vgat-IN | 0.400 | Kolmogorov-Smirnov |
| 1P | RF area; IC-PN/Vgat-IN | **<0.0001** | Kolmogorov-Smirnov |
| 1R | Spont. Rate; VPL-PN/IC-PN | **0.012** | Kolmogorov-Smirnov |
| 1R | Spont. Rate; VPL-PN/Vgat-IN | **<0.0001** | Kolmogorov-Smirnov |
| 1R | Spont. Rate; IC-PN/Vgat-IN | **0.0076** | Kolmogorov-Smirnov |
| 2C | Cdx2; 10 Hz Ctrl/+L | **0.0008** | Paired t-test |
| 2C | Cdx2; 50 Hz Ctrl/+L | **<0.0001** | Paired t-test |
| 2C | Cdx2; 300 Hz Ctrl/+L | **0.0013** | Paired t-test |
| 2E | Cdx2; Ind., Ctrl/+L | **0.0017** | Paired t-test |
| 2H | Adv; 10 Hz Ctrl/+L | **0.0003** | Paired t-test |
| 2H | Adv; 50 Hz Ctrl/+L | **<0.0001** | Paired t-test |
| 2H | Adv; 300 Hz Ctrl/+L | **0.0012** | Paired t-test |
| 2J | Adv; Ind. Ctrl/+L | **0.011** | Paired t-test |
| 2N | Surr. Inh., 10 Hz Ctr/+L | **0.0002** | Paired t-test |
| 2N | Surr. Inh., 50 Hz Ctr/+L | **<0.0001** | Paired t-test |
| 2N | Surr. Inh., Ind. Ctr/+L | **<0.0001** | Paired t-test |
| 3C | 10 Hz Ctrl/NBQX | 0.51 | Kolmogorov-Smirnov |
| 3C | 50 Hz Ctrl/NBQX | 0.16 | Kolmogorov-Smirnov |
| 3C | 300 Hz Ctrl/NBQX | 0.51 | Kolmogorov-Smirnov |
| 3L | Threshold Ctrl/NBQX | **0.0017** | Kolmogorov-Smirnov |
| 3M | 250 mN sus. Z-score Ctrl/NBQX | **0.0009** | Kolmogorov-Smirnov |
| 4L | RF Corr. Mech.-opt./Shuffled | **0.0003** | Kolmogorov-Smirnov |
| ED1 I,L | Conduction Velocity VPL/IC-PNs | 0.31 | Kolmogorov-Smirnov |
| ED2 P | RF area; VPL/IC-PN type 2 | **0.031** | Kolmogorov-Smirnov |
| ED3 D | PSDCs/LTMRs per segment | **0.017** | Paired t-test |
| ED5 F | Amplitude Ctrl./NBQX | **<0.0001** | Paired t-test |
| ED5 M | Threshold Ctrl/NBQX | 0.25 | Kolmogorov-Smirnov |
| ED5 M | 250 mN sus. Z-score Ctrl/NBQX | 0.76 | Kolmogorov-Smirnov |
| ED6 E | Behavior, total movement | **0.0047** | Kolmogorov-Smirnov |
| ED7 T | Threshold Ctrl/Lesion | **<0.0001** | Kolmogorov-Smirnov |
| ED7 U | 250 mN sus. Z-score Ctrl/NBQX | **<0.0001** | Kolmogorov-Smirnov |
| Text | DCN Spont. Rate Ctrl/DC Lesion | 0.93 | Kolmogorov-Smirnov |
| ED7 V | mVPL Spont. Rate Ctrl/DCN Lesion | **0.0014** | Kolmogorov-Smirnov |

## METHODS

### Animals

All experimental procedures were approved by the Harvard Medical School Institutional Care and Use Committee and were performed in compliance with the Guide for Animal Care and Use of Laboratory Animals. Animals were housed in a temperature- and humidity-controlled facility and were maintained on a 12:12 hour dark:light cycle. All experiments were performed on adult animals (>5 weeks) of both sexes. The following mouse lines were used: C57Bl/J6; *Calca-FlpE* (ref.^38^); *Avil^FlpO^* (ref.^38^); *Avil^Cre^* (ref.^57^); *Cdx2-Cre* (ref.^53^); *Rosa26^LSL-Acr^*^1^ (ref.^36^); *Rosa26^FSF-ReaChR^* (derived from ref.^58^); Animals were maintained on mixed C57Bl/J6 and 129S1/SvImJ backgrounds. C57Bl/J6 were obtained from Jackson Laboratories.

### Juxtacellular recordings

All recordings were performed in urethane-anesthetized mice. Adult (>5 weeks) animals were anesthetized with urethane (1.5 g/kg) and placed on a heating pad. The head and neck were shaved and local anesthesia (lidocaine HCl, 2%) was administered to the scalp and neck. An incision was made in the skin, and the muscle of the dorsal aspect of the neck was cut and moved aside to expose the brainstem. A headplate was attached to the skull using dental cement and the head was fixed to a custom-built frame. The dura overlying the brainstem was cut and small fragments of the occipital bone were removed. In some cases in which optical access to the DCN was required, small amounts of the posterior vermis of the cerebellum were aspirated to expose the DCN. A glass electrode filled with saline (2-3 MΩ) was put in place 200-300 µm above the gracile nucleus. The area was then flooded with low-melting point agarose dissolved in saline.

After hardening of the agarose, the electrode was advanced into the gracile. The electrode was guided to individual units and positioned to maximize the signal of a single unit. Recording quality and unit discrimination was continuously assessed using an audio monitor and online analysis of amplitude and spike waveforms. All recordings in the DCN were made in the hindlimb representation region of the gracile nucleus, and only units with receptive fields in the hindlimb were recorded. Recordings were targeted to the rostro-caudal level approximately where the gracile diverges bilaterally (see example in **Extended Data Fig. 1**) where units receiving input from the hindlimb digits and pads were most abundant. Although it has been reported that the gracile in rats, cats and primates is subdivided into ‘core’ and ‘shell’ regions, we were unable to detect clear organization in the mouse through electrophysiology experiments. Signals from single units were amplified using the 100x AC differential amplification mode of a Multiclamp 700B and sampled at 20 kHz using a Digidata 1550B controlled by Clampex 11 software (Molecular Devices). Signals were collected with an additional 20x gain, 0.3 kHz high pass filter, and a 3 kHz Bessel filter.

In some experiments DCN neurons were identified through antidromic activation of their axon in a target region. A craniotomy over the region of interest was performed. The head was leveled and bipolar electrodes (platinum-irridium 250 µm spacing, FHC) were lowered into the contralateral VPL of the thalamus (coordinates: 2 mm posterior to bregma, 2 mm lateral, 3.5 mm deep), or the contralateral inferior colliculus (coordinates: electrode angled 45 degrees, 4 mm posterior to bregma at tissue entry, 1 mm lateral, 1.7 mm deep). Single stimuli (60-200 µA) were applied every 3 sec while searching for units in the DCN. A collision test was performed for all units that could be antidromically activated. Stimuli were triggered from spontaneous spikes, and experiments were only continued for units that passed collision testing. We found that units with reliable and precisely timed antidromic spikes (jitter <1 ms) almost always passed a collision test. Units that failed collision testing had variable timing of activation and much longer latencies. Polysynaptic activation in the DCN was rarely detected by activating the inferior colliculus, and often observed when stimulating the VPL. In a subset of experiments, once data collection was complete, the stimulus electrode was retracted from the stimulation site and coated with DiI (ThermoFisher). The electrode was then advanced back into the tissue and left in place for at least 5 minutes. To verify that the electrode was in the same position as before, units were antidromically activated again. Electrodes were then retracted and the animal was perfused as described below for anatomical verification of the stimulus electrode position. In another subset of experiments, following identification of a VPL-PN, the stimulus electrode was then retracted from the VPL and moved to coordinates of the posterior nucleus of the thalamus (coordinates: 2 mm posterior to bregma, 1.1 mm lateral, 3 mm deep). Units that were originally antidromically activated from the VPL failed to be activated by stimulation in the new location (n/N: 3/3). Stimulation (60-200 µA) of the posterior nucleus also failed to evoke polysynaptic and multi-unit background activity in the DCN as was often seen with VPL stimulation.

To record from inhibitory neurons in the DCN, we used optical activation in *Vgat-ChR2* animals. Units were searched while applying 200-300 ms ramps of blue light (2-4 mW/mm^2^) from a fiber optic (400 µm diameter, 0.39 NA) placed above the DCN. Light was delivered from a 470 nm LED (M470F3, Thorlabs). Ramps were used because pulses of light were found to generate short latency activation in most units. Most units in the DCN are glutamatergic, and we found that pulsed light drives strong synchronized GABAergic input to primary afferent terminals and stimulates them through the depolarizing action of GABA, thereby driving vesicle release onto excitatory projection neurons. When ramps were used, many units were instead silenced, as expected. Units considered optotagged were those that could be activated during the light ramp.

### Optical silencing

For optical silencing of ascending inputs *in vivo*, experiments were prepared as described above except dissection was performed in the dark under red light illumination, as bright white dissection light was capable of activating Acr1. All experiments were performed in animals homozygous for the *Rosa26^LSL-Acr^*^1^ allele, as light was unable to fully silence inputs in heterozygous animals. Once the preparation was ready, the electrode was positioned in place above the DCN, along with a 400 µm diameter 0.39 NA fiber 1 mm above the DCN. The area was flooded with low-melting point agarose embedding the fiber and electrode. The electrode was then advanced into the DCN to record from single units.

Silencing trials were pseudo-randomly interleaved with control trials. On silencing trials, a 300- 400 ms light ramp (0.2-0.8 mW/mm^2^, 552 nm) was delivered to the surface of the DCN with mechanical stimuli delivered during the last 100-200 ms of the ramp. Light was delivered using a 552 nm LED (MINTF4, Thorlabs). We found that silencing became progressively ineffective after 500 ms of light delivery, possibly due to effects of prolonged depolarization of the terminals.

Optical silencing could only be performed successfully for gentle stimuli (≤ 20-30 mN). In *Cdx2*-Cre; *Rosa26*^LSL-Acr^^1^ animals, light was unable to fully silence inputs to the DCN when delivering intense indentations. Silencing using Acr may suppress vesicle release from ascending inputs substantially, but the simultaneous and ongoing activation of many inputs all at once may still allow postsynaptic DCN neurons to reach threshold despite a large reduction in the amount of synaptic drive. Thus, experiments were limited to gentle indentation and vibratory stimuli.

### Mechanical and optical stimuli

We selected for units that were primarily responsive to stimulation of the hindpaw. All mechanical stimuli were generated by a DC motor with an arm connected to a blunt and smoothed acrylic probe tip 1 mm in diameter. The use of a large diameter probe tip with smoothed edges allowed the delivery of high forces that that did not evoke paw withdraw in awake animals (**Extended Data Fig. 6**). We did not observe visible damage to the skin following high force stimuli in awake or anesthetized animals. The motor was driven by a custom-built current supply controlled by a data acquisition board (Digidata 1550B, Molecular Devices). Step indentation forces were calibrated using a fine scale. For experiments measuring responses to various forces, the probe tip was positioned in a resting state on the skin surface and indentations of incrementally increasing forces were applied every 3-10 seconds. Once the maximal force was reached, the force was reset to zero and was again incrementally increased. For experiments measuring responses to vibration, vibratory stimuli were applied at similar forces (10-20 mN) across frequencies. DCN units that did not respond to Pacinian-range vibration frequencies also did not respond when vibrations were delivered at higher forces.

The stimulator was attached to an articulating arm that could be moved by hand and would remain in position. For receptive field mapping, the probe tip was manually positioned over a single location of the hindlimb where it remained in place. A trial was then initiated to deliver a brief vibration (50 Hz, 100 ms, 10-20 mN). The experiment was performed under a stereoscope equipped with a CCD (BlackFly S BFS-U3-04S2C, Flir) that was triggered to capture images for each trial. This process was repeated until enough trials were acquired (60-100) to generate a receptive field map of the entire hindpaw. For experiments measuring indentation responses at various forces, the stimulator was firmly secured to a 0.5” heavy post in order to prevent relocation or repositioning when applying high forces.

Optical stimulation was performed on the skin or DCN using a 200 µm or 400 µm diameter 0.39 NA fiber coupled to a 554 nm LED (MINTF4, Thorlabs). For skin stimulation, the fiber tip was held in place with an articulating arm and moved in position manually for each trial, as performed for mechanical stimulation. For optical receptive field mapping, five pulses (2-5 ms duration, 60 mW/mm^2^) were delivered to the skin at 1 Hz for each trial.

### Multielectrode array recordings

Recordings in the thalamus were made in the middle VPL (2 mm bregma, 2 mm lateral, 3.5 mm deep). Animals were headplated, the DCN was exposed as described above, and a craniotomy above the VPL was performed. The dura was removed, and the area was flooded with 2% low-melting agarose dissolved in saline. A 32-channel multi-electrode array (MEA, A1x32-poly2- 10mm-50s-177-A32, Neuronexus) was lowered at 5 µm/s into the brain. Once positioned in a region where firing in many units could be evoked by brushing the hindpaw, the MEA was kept in place for 20 minutes to ensure a stable recording. The hindpaw was embedded in modeling clay for stabilization and the receptive fields of units were quickly assessed using a brush. The mechanical stimulator probe tip was then placed over the region of the hindpaw that could maximally activate the most units. Step indentations of 0-300 mN were applied every 3-6 seconds in ascending order and repeated until at least 10 trials per force were obtained.

Following stimulation the DCN was lesioned, either by using a 30 gauge needle and striking through the gracile nucleus, or by aspirating the gracile nucleus. Step indentations were then repeated. Throughout the experiment, the waveform of a single unit was monitored closely to assess drift, and any experiment with detectable waveform changes were discarded. In some cases, once the experiment was complete, the MEA was retracted from the brain, coated with DiI, and then descended to the same coordinates and allowed to stabilize for 5 minutes. Animals were then anesthetized with isoflurane and transcardially perfused with PBS followed by 4% PFA in order to assess the location of the lesion and electrode placement.

For lesions of the DC, animals were prepared as described above, and a laminectomy was performed at approximately T12 vertebrae and the dura was removed. A high-density 32-channel MEA (A1x32-poly3-5mm-25s-177-A32, Neuronexus) was inserted into the gracile nucleus and allowed to stabilize for 15 minutes. A baseline of 5 minutes of spontaneous firing was collected. The DC was then lesioned at approximately T12 using a 30 G needle. Brief vibratory stimuli were applied to the hindlimb throughout the experiment to assess the effectiveness of the lesion.

MEA recordings were made using an Intan headstage, recording controller (Intan Technologies RHD2132 and Recording Controller) and open-source acquisition software (Intan Technologies RHX data acquisition software, version 3.0.4). Data was sampled at 20 kHz and bandpassed (0.1 Hz – 7.5 kHz).

### *Ex vivo* recordings

Intracellular recordings were performed in random DCN neurons ex vivo. The brainstem was prepared as previously described^13^, except DCN neurons were recorded directly from the dorsal surface of a dissected brainstem. Borosillicate electrodes (2-3 MΩ) filled with internal solution consisting of in mM: 130 K-gluconate, 3 KCl, 10 HEPES, 0.5 EGTA, 3 MgATP, 0.5 NaGTP, 5 Phosphocreatine-Tris_2_, 5 Phosphocreatine-Na_2_, pH 7.2 with KOH, were visually guided to the DCN. Experiments were performed in recirculated ACSF containing in mM: 127 NaCl, 2.5 KCl, 1.25 NaH_2_PO_4_, 1.5 CaCl_2_, 1 MgCl_2_, 26 NaHCO_3_, 25 Glucose. Oxygenated with 95% O_2_ / 5% CO_2_. The preparation was kept at 35°C using an in-line heater. Recordings of random DCN cells were made using a Multiclamp 700B (Molecular Devices) and acquired using a Digidata 1550B with Clampex 11 software (Molecular Devices).

### Pharmacology

For spinal cord silencing experiments, animals were prepared for juxtacellular DCN recordings as described above. A laminectomy was performed over L3-5 spinal segments, the region of the spinal cord responsive to hindlimb stimulation. For control experiments, a laminectomy was performed over T10-T13 spinal segments. The exposed spinal cord and vertebral column was then flooded with low-melting point agarose. Once hardened, agarose overlying the dorsal horn was cut away to create a pool and confine drugs to the spinal cord. The dura was removed from the spinal cord using fine forceps. Prior to drug application, stimuli were applied to the hindpaw to evoke typical responses to ensure that the spinal cord or dorsal column had not been damaged. Drugs were then applied to the spinal cord. The non-competitive NMDAR antagonist MK-801 (10 mM, 10 µL, Abcam; dissolved in 90% H_2_O, 10% DMSO) was applied to the surface of the spinal cord and allowed to enter the cord for 3-5 minutes. The surface of the cord was then irrigated with saline. NBQX (10 mM, 20 µL, Abcam, dissolved in H_2_O) was then applied to the surface of the cord. After 5 minutes, the cord was covered with gelfoam which was allowed to absorb the NBQX and remained in place for the duration of the experiment, occasionally re-wet with saline. Drugs were applied sequentially in order to prevent precipitates from drug mixing at high concentration. Units were then recorded, and animals were sacrificed within 2-3 hours following drug application.

For controls measuring the efficacy of blockade, a laminectomy was performed over the L4 spinal cord. The vertebral column was held in place by two custom spinal clamps. The dura was removed using fine forceps or a fine needle and kept moist with saline. A 32-channel MEA was inserted into medial L4 (A1x32-poly2-10mm-50s-177-A32, Neuronexus). Step indentations (300 mN) were applied at 0.1 Hz, and the MEA was lowered such that the most dorsal channel detected minimal evoked responses. The location of the dorsal channel was assumed to be near the surface of the cord. The MEA was then kept in place for 15 minutes. Baseline trials were collected and then drugs were applied as described above.

### Stereotaxic injections

Adult animals were anesthetized with isoflurane and placed in a stereotaxic frame. The head was tilted 30° forward. The hair over the neck and caudal scalp were removed using a clipper and the skin was sanitized using isopropanol followed by betadine. Local anesthesia (2% lidocaine HCl) was applied to the area and an incision was made to expose neck muscles. Neck muscles were separated from the skull to expose the brainstem. A 30-gauge needle was used to cut the dura overlying the brainstem and expose the DCN. A pipette filled with retrograde tracer (Red Retrobeads, Lumafluor, or cholera toxin subunit B 2 µg/µL, Fisher) and 0.01% fast green was lowered into the DCN just sufficient to penetrate the surface. Once penetrating the surface, a small volume (30-50 µL) of retrograde tracer was injected. The pipette was held in place for 20 seconds and then removed. Care was taken to ensure tracer did not leak from the injection site following injection, and that tracer labeled with fast green filled the DCN, but did not extend beyond the nucleus. The pipette was then removed, and overlying muscle and skin was sutured shut. Animals were administered analgesic (Buprenex SR, 0.1 mg/kg, ZooPharm) prior to surgery and monitored post-operatively. Following 1-3 days, animals were transcardially perfused for tissue harvest.

### Histology

Animals were anesthetized with isoflurane and transcardially perfused with PBS followed by 4% PFA in PBS. Brains were removed and post-fixed in PFA overnight. The brain, spinal cord and DRGs were dissected free. The isolated spinal cord or brain was mounted in low-melting point agarose and the thoracic spinal cord 60 µm thick sections were made on a Leica VT1000S vibratome. For immunohistochemistry, free-floating sections were first permeabilized with 0.1% Triton-X100 in PBS for 30 min at room temperature. Sections were then incubated with 0.1% Triton-X100 and 4% normal goat serum (NGS, Abcam) for 30 min at room temperature. Primary antibodies were then added (mouse anti-NeuN, 1:1000, MAB377, Millipore; guinea-pig anti- Vglut1, 1:2000, Synaptic Systems 135302) and incubated overnight at 4°C. Sections were then washed three times with PBS with 0.1% Triton-X100 and 4% NGS for 10 minutes at room temperature. Secondary antibodies were then applied (IB4-Alexa 647, 1:300, ThermoFisher I32450; goat anti-mouse Alexa-488, 1:500, Abcam ab150113; FITC goat anti-GFP, Abcam ab6662) for 2 hours at room temperature, and washed with PBS three times for 10 minutes.

Sections were mounted onto glass slides using Fluoromount Aqueous mounting medium (Sigma). Sections were imaged with a Zeiss LSM 700 confocal microscope using a 20x 0.8 NA oil immersion objective using Zen software. DRGs were whole-mounted and imaged using a Zeiss LSM 700 confocal microscope using a 10x 0.45 NA air objective.

PSDCs were counted in Z-stacks (3-4 µm Z-spacing) of 60 µm thick thoracic spinal cord sections. 10-20 sections were analyzed per animal, and the average number of PSDCs per section was measured and multiplied by 16.67 to estimate the number PSDCs in one segment (1000 µm) of spinal cord for that animal. Aβ-LTMRs were counted in whole-mounted thoracic DRGs and compared to spinal cords of the same animals. Labeled DRG neurons were counted in Z-stacks of the entire DRG (5-6 µm Z-spacing). If more than one DRG was analyzed per animal, the number of counted cells was averaged between the two.

### Behavior

Behavior was performed in two C57Bl/J6 males and four *Calca-FlpE*; *Rosa26^FSF-ReaChR^* animals (2 male, 2 female). Animals were anesthetized with 2% isoflurane. Hair over the scalp was clipped using a shaver and skin was disinfected using isopropanol followed by betadine. Local anesthesia (2% lidocaine HCl) was applied to the scalp. An incision was made and the dorsal skull was cleared of skin and muscle using a scalpel. A headplate was rested on top of the skull and fixed in place using dental cement. Animals were administered analgesic (Buprenex SR, 0.1 mg/kg, ZooPharm) prior to surgery and monitored post-operatively.

Animals were allowed to recover for 2-3 weeks. Headplated animals were then transferred to a rig consisting of an acrylic platform. Animals were headfixed to a suspended post, but were otherwise free to move. Mice were allowed to habituate to the rig for 5-10 minutes. Gentle mechanical stimuli were then delivered to prevent startle responses. First a brush was used to gently stroke the trunk and hindlimbs. After 5-10 strokes with a brush, the indenter was introduced and 300 ms indentations of 10 mN were delivered using the same smoothed 1 mm probe tip used in electrophysiology experiments. The indenter and probe tip were the same as those used for experiments in anesthetized animals. Indentations were delivered to the middle of dorsal hindpaw as the ventral side was inaccessible in awake animals with unrestrained paws.

When the animal was still, the indenter tip was positioned in place above the hindpaw, a trial was triggered and indentation was delivered after one second of baseline. Stimuli were delivered approximately every 20 seconds. After a few trials using 10 mN stimuli, animals no longer were startled by indentation, and the force was increased to 300 mN. Trials were recorded using a CCD (BlackFly S BFS-U3-04S2C, Flir) attached to a stereoscope with frames captured at 100 Hz. The first trial of 300 mN typically evoked a startle response, and was discarded. The following trials were then collected for analysis (5-12 trials). Several seconds following 300 mN indentation, a small force (<10 mN) was briefly applied to the probe in order to bring it into position near the skin again for the subsequent trials.

In four *Calca*-*FlpE*; *Rosa26^FSF-ReaChR^* animals, once mechanical stimulation was completed, a set of trials were performed in which the skin was optically stimulated. A 400 µm 0.39 NA fiber coupled to a 554 nm LED was held in place above the center of the dorsal hindpaw. A 50 ms pulse of light (∼30 mW/mm^2^) was delivered to the skin. The first trial was discarded, and subsequently 4-7 trials were collected for analysis. Experiments were ended after fewer trials because animals showed clear signs of pain.

In order to determine whether mechanical stimulation generated enough force to prevent the animals from withdrawing their paw, optical stimulation was delivered during mechanical stimulation in two *Calca*-*FlpE*; *Rosa26^FSF-ReaChR^* animals. A 300 mN, 300 ms indentation was delivered as described above. 100 ms after the onset of indentation, a 50 ms pulse of light was delivered at or near the indentation as described above. Animals readily withdrew their hindlimb within 30 ms of light onset when 300 mN of force was still being applied.

### Analysis

Juxtacellular recordings in the DCN were analyzed offline using custom written scripts in Matlab (Mathworks). Spikes were detected using an amplitude threshold and recordings in which the unit could not be isolated by amplitude were discarded.

Receptive field maps were generated using custom written scripts in Matlab. An image of the hindpaw from the experiment was used as a template. Images capturing the probe or fiber optic for each trial were cycled through and the location of the probe or fiber tip was manually marked on the template image. The coordinates of each stimulus location and the template image were then combined with electrophysiology data to identify the number of evoked spikes for each stimulus location. Each unit’s receptive field map is displayed with points indicating the location of the stimulus and normalized change in firing rate from baseline. The maximal change in firing rate is indicated in text within the image. Maximal firing rate was measured as the number of spikes over the course of the 100 ms vibratory stimulus, or number of spikes within a 20 ms window following optical stimulation. The spontaneous firing rate was measured as the average firing rate prior to mechanical stimulation across trials. In order to measure the receptive field area, a threshold was taken at the half-maximal evoked firing rate. Stimulus locations that evoked the half-maximal firing rate or more were included in the receptive field. The area bound by these points was used as a measure of the receptive field area.

Correlation of optical and receptive fields were performed by comparing regions of the hindpaw. First, the optical and mechanical receptive fields were mapped as described above. The paw was then subdivided into segments corresponding to digits and pads. The stimulus trials falling within each segment were averaged to obtain an average evoked firing rate for each segment. The optically-evoked firing rate for each segment was plotted against its mechanically evoked firing rate for that same region. This generated a plot of 17 points, each representing the optical and mechanical response of one region of the hindpaw. A linear fit was performed and the Pearsons correlation coefficient (R^2^) was used as a measure of receptive field correlation. As a control, similar analysis was performed but optical and mechanical receptive fields were shuffled between units.

### Spike sorting (MEAs)

MEA recordings underwent initial analysis using Kilosort 2.0 (ref. ^59^). Prior to analysis, baseline and lesions trials were interleaved in order to prevent artifactual drift corrections by Kilosort.

Drift monitoring was performed during acquisition and experiments with detectable changes in spike waveforms were discarded. Default detection settings were used for analysis, except template amplitude thresholds were set to [5 2]. Clustered were manually curated using Phy^60^ by examining spike waveforms and autocorrelations in order to identify putative single units. As receptive fields were not mapped for MEA experiments, units underwent selection for inclusion. Units that had low thresholds at baseline (<10 mN), or units that had mechanically-evoked firing that was unaffected by lesioning the DCN were included. Units that had high-thresholds (>10 mN) at baseline and became mechanically insensitive following DCN lesion were not included for analysis because the stimulus may have been off the center of the receptive field.

Z-scores were computed by measuring the average baseline firing rate within a 0.5-1 s window prior to stimulation. For optical silencing, pharmacology and lesion experiments in which the same units were monitored before and after manipulation, Z-scores in experimental conditions were computed from the mean and standard deviations of baseline trials.

### Behavior

Videos of mechanical and optical stimuli delivered to the hindpaw were collected in trials lasting 3 seconds. The paw position was semi-automatically tracked using the Video Labeler app in Matlab. Baseline paw position was collected 0.5 s prior to stimulus delivery for the trial. The baseline position was set as the origin, and position of the paw relative to the starting position was measured for the entire trial. For average plots, the position was averaged across trials for each animal. For total movement, the total change in position was summed over the average trial for each animal.

### Statistical analyses

Statistical analyses were performed with significance threshold set at p < 0.05. All summary data is presented as mean ± SEM unless otherwise noted. All t-tests were two-tailed.

### Data availability

Data generated during this study will be included in this published article (and its supplementary information files) upon acceptance of the manuscript.

### Code availability

Custom scripts used in this study will be posted to Github upon acceptance of the manuscript.

## ACKNOWLEDGEMENTS

We thank Christopher Harvey, Chinfei Chen, Wade Regehr, Celine Santiago, Alan Emanuel, Genelle Rankin, Andrew Shuster, Anda Chirila, Rosa Martinez-Garcia, Dawei Zhang, and Jessica Barowski for comments on the manuscript. This work was supported by a Mahoney Postdoctoral Fellowship (JT), a Gordon Postdoctoral Fellowship (JT), NIH grant NS097344 (DDG), The Hock E. Tan and Lisa Yang Center for Autism Research at Harvard University (DDG), and the Edward R. and Anne G. Lefler Center for Neurodegenerative Disorders (DDG). DDG is an investigator of the Howard Hughes Medical Institute. This article is subject to HHMI’s Open Access to Publications policy. HHMI lab heads have previously granted a nonexclusive CC BY 4.0 license to the public and a sublicensable license to HHMI in their research articles. Pursuant to those licenses, the author-accepted manuscript of this article can be made freely available under a CC BY 4.0 license immediately upon publication.

## AUTHOR CONTRIBUTIONS

JT, BPL, and DDG conceived the project. JT performed all experiments except data shown in Extended Data Fig. 3. JT performed all analysis. BPL performed pilot experiments not presented, and anatomical experiments shown in Extended Data Fig. 3. JT and DDG wrote the paper with input from BPL.

## COMPETING INTERESTS

The authors declare no competing interests.

## Notes

### Competing Interest Statement

The authors have declared no competing interest.

### Summary of Updates

Figures 2,4, and Extended Data Figures 1-2,4-9 have been updated with new additional experiments.

